# Genetic sex of enteric neurons enables ovarian relaxin to gate maternal gut plasticity

**DOI:** 10.64898/2026.01.09.698651

**Authors:** Bryon Silva, Edisona Tsakani, Alexandros Kontopoulos, Joydeep De, Pedro Gaspar, Alya Hiridjee, Camila Rosselli, Jaime de Juan-Sanz, Jorge M Campusano, Christoph D. Treiber, Scott Waddell, Irene Miguel-Aliaga, Dafni Hadjieconomou

**Affiliations:** The Francis Crick Institute, London, UK; MRC Laboratory of Medical Sciences, London, UK; Institute of Clinical Sciences, Faculty of Medicine, Imperial College London, UK; Institut du Cerveau-Paris Brain Institute (ICM), Sorbonne Université, Inserm, CNRS, Hôpital Pitié-Salpêtrière, Paris, France; Facultad de Ciencias Biológicas, Pontificia Universidad Católica de Chile, Santiago, Chile; Department of Biology, University of Oxford, UK; Centre for Neural Circuits and Behaviour, University of Oxford, UK

## Abstract

Animals must align intestinal plasticity and feeding with reproductive state, yet the checkpoint that gates these adaptations is unknown. Here we show that an ovary-to-enteric-neuron axis gates the onset of maternal gut plasticity in Drosophila. Genetic sex establishes endocrine competence in a subset of enteric neurons via the sex determination pathway, enabling female-specific expression of the relaxin-family receptor Lgr3. After mating, steroid signalling increases Lgr3 receptor expression, priming these neurons for reproductive adaptation. Once oocytes mature fully, follicle cells secrete the relaxin-like hormone dILP8, which activates Lgr3 to trigger gut enlargement and increased feeding. Disrupting the sex determination pathway in enteric neurons, Lgr3, or ovarian dILP8 prevents gut enlargement and reduces feeding. Thus, genetic sex establishes competence, steroid signalling primes it, and ovarian relaxin triggers it, defining a maternal intestinal plasticity checkpoint that ensures gut adaptations initiate only once reproduction is underway and energy demands peak. Our findings delineate an ovary-to-enteric-neuron axis that couples reproductive state to maternal gut plasticity.

## Main

Life-stage transitions reprogramme organismal metabolism^1–3^. In mammals, reproduction reprogrammes hypothalamic circuits to enable maternal behaviours^4–7^, establishing that reproductive state can reconfigure neuronal function. Across species, reproduction in females enlarges the gut and increases feeding^1,8^, yet how these intestinal adaptations are initiated is unknown. The enteric nervous system (ENS), which coordinates gastrointestinal function and integrates endocrine signals, links reproductive state to changes in intestinal circuit activity^1,9^. Recent findings support roles for systemic steroid hormones and gut-derived peptides in reproductive intestinal remodelling^1,10,11^, but the mechanisms by which gonadal state is relayed to enteric circuits to orchestrate intestinal plasticity have not been described. Identifying such a mechanism would reveal how reproductive status aligns intestinal plasticity with the energetic demands of reproduction.

## Results

### The genetic sex of enteric Ms neurons establishes competence for maternal gut plasticity

To investigate mechanisms that enable enteric neurons to respond to reproductive cues, we first asked whether enteric neurons have a cell-intrinsic sexual identity, a feature that could influence their ability to respond to sex-matched reproductive cues. In somatic and gonadal tissues alike, genetic sex is determined by the activity of the sex determination pathway^12–15^. Sexual identity maintained into adulthood can shape tissue-specific physiological plasticity^16–21^. This principle has been well illustrated in contexts such as gut epithelial renewal and proliferation^11,16^. However, how genetic sex impinges on the function of gut-innervating neurons remains unknown across the evolutionary tree.

We focused on a specific subset of enteric neurons, the Myosuppressin (Ms) expressing neurons, which innervate the crop (a gut compartment functionally analogous to the mammalian stomach^9^) and mediate post-mating gut plasticity^1^. We queried the involvement of the canonical sex determination pathway by using the *transformer (tra)* gene, which is transcribed in both sexes but undergoes female-specific splicing to produce a functional Tra protein only in females^22^ (**Extended Data Fig. 1a)**, and governs sex-specific transcriptional programmes^23^. An endogenous knock-in reporter of *tra*^24^ drove expression in Ms neurons of adult female flies but not in those of males, supporting the idea that Ms neurons are intrinsically sexually dimorphic (**Fig. 1a, Extended Data Fig. 2a**). Thus, Ms neurons exhibit a female-specific identity established by *tra*, positioning them as a potential site where female reproductive signals converge with enteric physiology.

**Figure 1.**
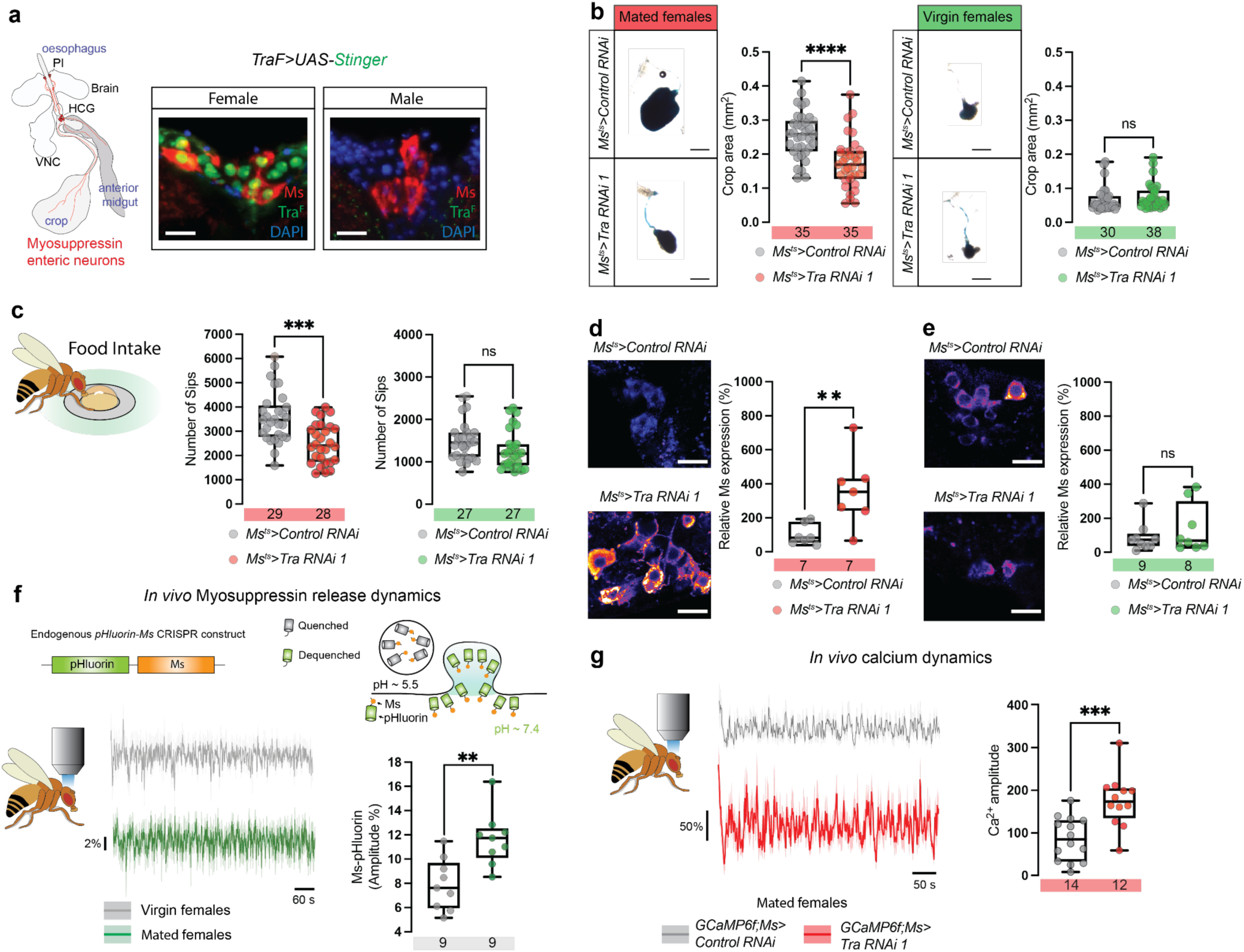
Genetic sex of Ms enteric neurons controls crop plasticity, food intake, and neuronal activity in mated females. **(a)** Left, schematic of the *Drosophila* digestive tract highlighting Ms neurons (red) innervating the crop. The somata of these gut neurons reside both in the pars intercerebralis (PI) and hypocerebral ganglion (HCG). From these two centres of visceral innervation, Ms neurons project via the recurrent nerve to the crop and anterior midgut^1^. Right, confocal images of Ms neurons in *TraF^KI^>UAS-Stinger* flies show nuclear GFP (green) and Ms peptide (red) expression in females, but not in males, indicating sex-specific expression of *TraF*. Scale bar: 10 μm. **(b)** Representative images (top) and quantification (bottom) of crop size in mated females (red), and virgin females (green) expressing either control RNAi or *Tra* RNAi under *Ms*-Gal4. *Tra* knockdown significantly reduces crop area in mated females (****p < 0.0001), but has no effect in virgins (ns: not significant). n values are indicated on each plot. Scale bar: 200 μm. **(c)** Quantification of food intake (number of sips) in mated females and virgin females using the FlyPAD assay. *Tra* knockdown in Ms neurons significantly reduces food intake in mated females (***p < 0.001), but not in virgin females. **(d–e)** Confocal images and quantification of *Ms* protein expression in Ms enteric neurons of mated (d) and virgin (e) females. *Tra* knockdown in Ms neurons increases *Ms* expression in mated females (**p < 0.05), but not in virgins (ns). Scale bars: 10 μm. **(f)** Endogenous Ms-pHluorin shows larger exocytic events after mating (**p<0.01). Schematic of the CRISPR Ms-pHluorin knock-in allele and pH-dependent dequenching. Average of individual traces from neuronal somas of virgin females (grey) and mated females (green). Right, amplitude (%), each dot represents one fly brain. **(g)** Calcium imaging of Ms neurons using GCaMP6f in mated females, showing the average traces (left) and quantification of Ca²⁺ amplitude (right). *Tra* knockdown significantly increases calcium dynamics amplitude in Ms neurons (***p < 0.001). n values are shown on each plot.

Gut-innervating Ms neurons release Ms that activates the MsR1 receptor in the crop muscles^1^, thereby enabling crop expansion to increase food intake. This post-mating crop enlargement is required to meet the increased energetic demands of reproduction^1^. We therefore tested whether the female-specific identity determined by *tra* expression is required for this gut plasticity. To test this, we masculinised^16,25^ Ms neurons by downregulating *tra* specifically in adult Ms neurons using *UAS-tra RNAi* under the control of the *Ms-Gal4* driver^1,26^ in combination with the ubiquitously expressed temperature-sensitive GAL4 repressor, GAL80^27^, thereby restricting expression to adulthood and preventing developmental perturbations (see Methods for details). Masculinisation of Ms neurons significantly reduced crop expandability in mated females (**Fig. 1b**) but had no effect in virgin females. Similar results were obtained using a second, non-overlapping RNAi construct (**Extended Data Fig. 2b**). Therefore, *tra* is necessary in Ms neurons of female flies to drive the crop enlargement that occurs after mating.

We next tested whether *tra* in Ms neurons also regulates food intake, a second key component of reproductive metabolic adaptation^1^. Strikingly, by masculinising Ms neurons in adult mated females, they failed to show the typical post-mating increase in food intake (**Fig. 1c**), while feeding in virgin females remained unaffected. These data suggest that *tra* is essential for Ms neurons to coordinate the behavioural and physiological competence required to meet reproductive nutritional needs.

Since crop expandability and increased food intake are both mediated by the release of the Ms neuropeptide^1^, we hypothesised that *tra* may regulate Ms neuronal output. Indeed, we found that the downregulation of *tra* caused a marked accumulation of Ms neuropeptide in Ms neurons of mated females (**Fig. 1d, Extended Data Fig. 2c**), that was consistent with a lack of peptide release upon mating^28^. This effect was not observed in virgin females (**Fig. 1e, Extended Data Fig. 2d**), suggesting that *tra* controls Ms neuronal plasticity specifically in response to reproductive cues. To directly monitor Ms secretion *in vivo*, we CRISPR-engineered an endogenous pHluorin-Ms knock-in (see Methods), fusing pHluorin to the Ms peptide at the native locus (**Fig. 1f**). pHluorin is quenched in the acidic lumen of dense-core vesicles and dequenches upon exocytosis, yielding a transient fluorescence spike that decays as vesicles re-acidify^29^. Using cultured rodent neurons, we found that Ms-pHluorin robustly reported an increase in the number of exocytosis events during field stimulation *in vitro* (**Extended Data Fig. 2e**). Similarly, *in vivo* pHluorin-Ms imaging of Ms enteric neurons revealed larger transient amplitudes in mated compared to virgin females (**Fig. 1f**), consistent with a greater probability of exocytic events.

To further examine the function of Ms neurons, we performed *in vivo* calcium imaging of Ms neurons in mated females. We have previously shown that Ms neurons of mated females exhibit reduced amplitude of calcium oscillations compared to virgin females^1^. Downregulation of *tra* in Ms neurons increased the amplitude of *in vivo* calcium oscillations in the Ms neuron cell bodies of mated females (**Fig. 1g**), similarly to those observed in virgin females^1^. Together, these findings show that the sex determination gene *tra* establishes a female-specific programme in enteric neurons, enabling them to activate the neuroendocrine responses required for reproductive plasticity. This sexually dimorphic identity ensures that, upon mating, Ms neurons can initiate the female crop adaptations needed to support reproduction.

### The relaxin receptor Lgr3 is required for reproductive plasticity in enteric neurons

Having established that Ms neurons acquire a female-specific identity essential for reproductive gut plasticity, we next investigated how this identity is translated into functional output. To this end, we sought molecular effectors that might act downstream of *tra* to mediate the reproductive role of Ms neurons. We analysed a previously published single-cell RNA sequencing dataset^30^ and performed a pseudobulk differential expression analysis directly comparing female and male Ms neurons, identifying sex-specific molecular signatures in Ms neurons (see Methods for details). Among the transcripts enriched in female Ms neurons, we identified *Lgr3*, a relaxin-family G protein–coupled receptor^31^ that showed strong female-biased expression (higher fraction of expressing cells and higher mean expression), with lower detection in males (**Fig. 2a, Extended Data Fig. 1b**). The female-specific expression of Lgr3 in Ms neurons led us to hypothesise a role in female reproductive physiology.

**Figure 2.**
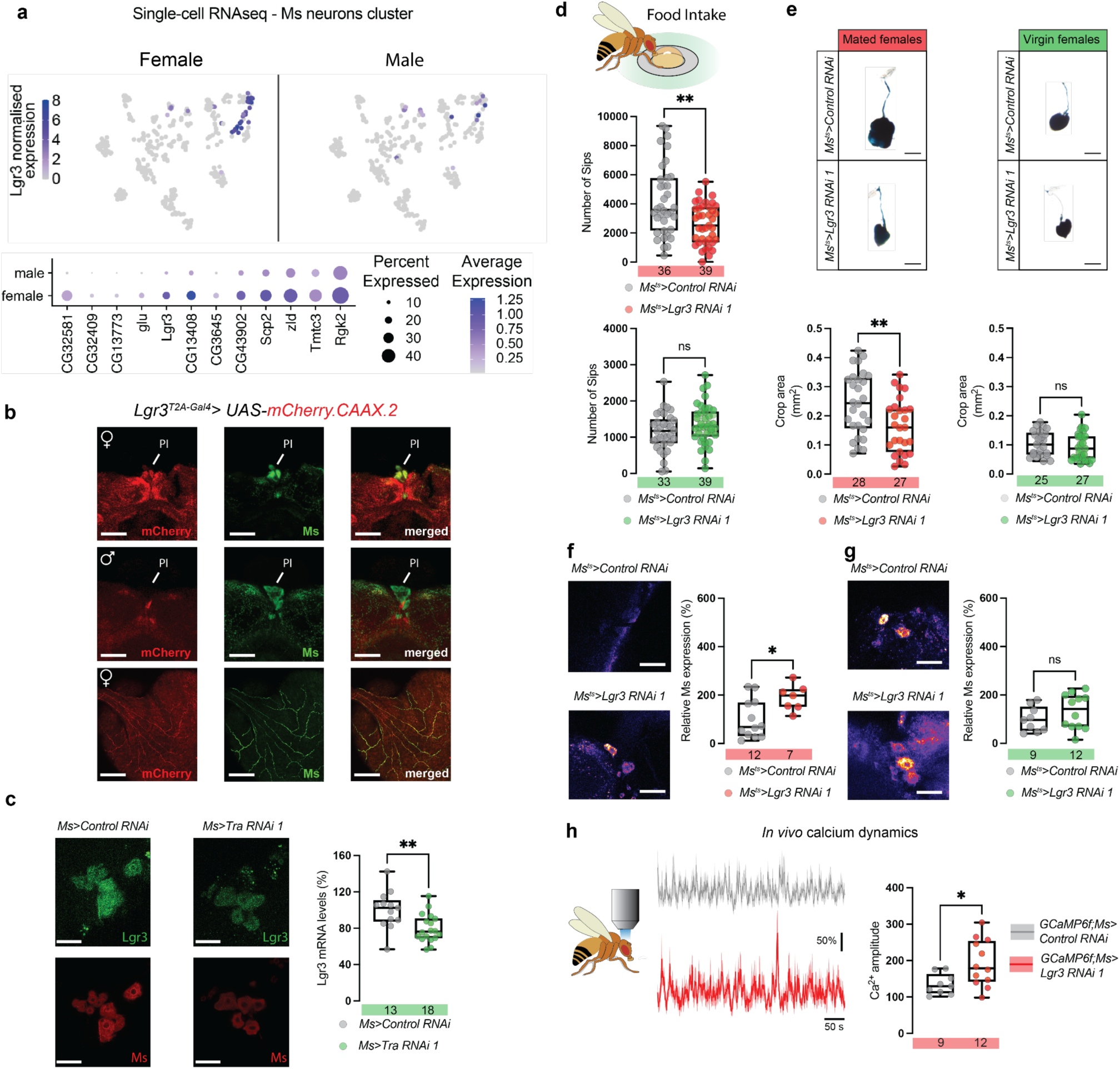
Relaxin receptor Lgr3 is a female-specific effector downstream of *transformer* in Ms neurons controlling crop plasticity, food intake and neural activity. **(a)** Single-cell RNA-seq analysis reveals *Lgr3* as a female-enriched receptor in Ms neurons. Analysis of a published adult brain scRNA-seq dataset (See Methods) was used to identify Ms-neuron clusters. Top: UMAP embedding of Ms-positive neurons from an adult brain scRNA-seq dataset, shown separately for females (left) and males (right); each dot represents a single Ms-positive neuron coloured by normalised Lgr3 expression. Bottom: dot plot summarising selected genes from a pseudobulk differential expression analysis directly comparing female and male Ms neurons; dot size indicates the fraction of Ms cells expressing each gene and colour denotes average expression. **(b)** Confocal images showing female-specific co-expression of *Lgr3* in Ms neurons using *Lgr3T2A-Gal4>UAS-mCherry-CAAX.2*. Ms neurons (in green), *Lgr3*-driven membrane mCherry (in red), and merge confirms co-localisation. In females, Lgr3 is present in Ms somata and co-localises with Ms. In males, reporter signal is not detected in Ms neurons under identical acquisition settings. Bottom row, magnification view of Ms projections in the crop terminals. Scale bars: 10 μm (top and middle), 200 μm (bottom). **(c)** In situ hybridization for *Lgr3* mRNA (green) in Ms neurons (red) shows reduced *Lgr3* transcript levels in *Ms*>Tra RNAi virgin females compared to controls. Quantification confirms significant reduction (**p < 0.01). Scale bar: 10 μm. **(d)** Food intake measured by FlyPAD in mated females, and virgin females. *Lgr3* knockdown in Ms neurons significantly reduces sip number in mated females (**p < 0.01), but not in virgin females. **(e)** Representative crop images and quantification of crop size across conditions. Knockdown of *Lgr3* in Ms neurons significantly reduces crop size in mated females (**p < 0.01), but has no effect in virgin females (ns: not significant). Scale bars: 200 μm. **(f–g)** Confocal images of *Ms* peptide levels in Ms enteric neurons of mated (f) and virgin (g) females. *Lgr3* knockdown increases *Ms* peptide levels in mated females (*p < 0.05) but not in virgins (ns). Scale bars: 10 μm. **(h)** Calcium imaging average traces (left) and quantification of Ca²⁺ amplitude (right) in Ms neurons using GCaMP6f. *Lgr3* knockdown increases the amplitude of spontaneous calcium dynamics in mated females (*p < 0.05). n values are indicated in the boxplots.

To verify this finding, we examined Lgr3 expression using two independent reporter lines: a transcriptional *Lgr3-T2A-Gal4^CR^*^00347^ reporter^32^ and an endogenous N-terminal GFP-tagged Lgr3 (*sfGFP::Lgr3*) protein reporter^33^. Both reporters confirmed expression of *Lgr3* in Ms neurons of virgin females (**Fig. 2b, Extended Data Fig. 3a-g, Extended Data Fig. 4a,b**), whereas male Ms neurons showed low and infrequent reporter signal (only ∼25% of the examined brains contained at least one Ms⁺Lgr3⁺ neuron), consistent with the single-cell RNA-seq female bias expression. Since *tra* is expressed in female Ms neurons, we asked whether *tra* regulates *Lgr3* expression. Single-molecule RNA fluorescence *in situ* hybridisation (smRNA FISH, see Methods for details) showed that Ms-specific *tra* knockdown significantly reduced Lgr3 mRNA in virgin females (**Fig. 2c**). This indicates that *tra* is not only necessary for the female identity of Ms neurons, but also supports the expression of *Lgr3* in virgin females. These findings suggest that *tra* confers the female genetic sex on Ms enteric neurons, at least in part, by regulating the expression of the relaxin receptor *Lgr3*.

We next asked whether *Lgr3* is functionally required in Ms neurons to drive reproductive gut plasticity. Knockdown of *Lgr3* in Ms neurons significantly impaired food intake and crop expandability in mated females but had no effect in virgin females (**Fig. 2d,e**), consistent with an inability to engage reproductive feeding programmes. These effects were confirmed using a second RNAi construct targeting a non-overlapping region of *Lgr3* (**Extended Data Fig. 5**).

Since Ms neurons regulate crop expandability and feeding via neuropeptide release, we next asked whether *Lgr3* is required for this output. *Lgr3* knockdown led to marked accumulation of Ms peptide in mated females (**Fig. 2f, Extended Data Fig. 4c**), while peptide levels in virgin females (**Fig. 2g, Extended Data Fig. 2d**) remained unchanged, suggesting that Lgr3 is required for Ms peptide release from the Ms neurons of mated females that would deplete intracellular Ms peptide stores. These data indicate that *Lgr3* is required for mating-dependent activation of Ms neurons.

We next examined the function of Ms neurons via *in vivo* calcium imaging in mated females. Downregulation of *Lgr3* in Ms neurons increased the amplitude of *in vivo* calcium oscillations of mated females (**Fig. 2h**), further supporting a role of Lgr3 function in Ms neurons.

Altogether, our findings demonstrate that Lgr3 expression in female Ms neurons is essential for function. Tra defines the female neuronal genetic sex of Ms neurons and supports their female-enriched Lgr3 expression, thereby granting endocrine competence to respond to a reproductive signal that coordinates gut plasticity with reproductive demands.

### Steroid hormone signalling primes Ms neurons after mating

Having shown that the relaxin receptor Lgr3 is required in Ms neurons for reproductive gut plasticity, we next asked whether its expression is modulated by reproductive state. In non-reproductive tissues, reproductive cues can dynamically reshape gene expression to synchronise tissue physiology with reproductive demands^34,35^. To test whether this applies to Lgr3 in enteric neurons, we quantified its expression in Ms neurons of virgin and mated females. We found that both *Lgr3* mRNA and protein levels were significantly higher in Ms neurons of mated females compared to virgin females (**Fig. 3a,b, Extended Data Fig. 6a**), indicating that mating induces Lgr3 upregulation. This post-mating increase in Lgr3 suggested the involvement of a potential systemic hormonal signal that primes Ms neurons for reproductive adaptation.

**Figure 3.**
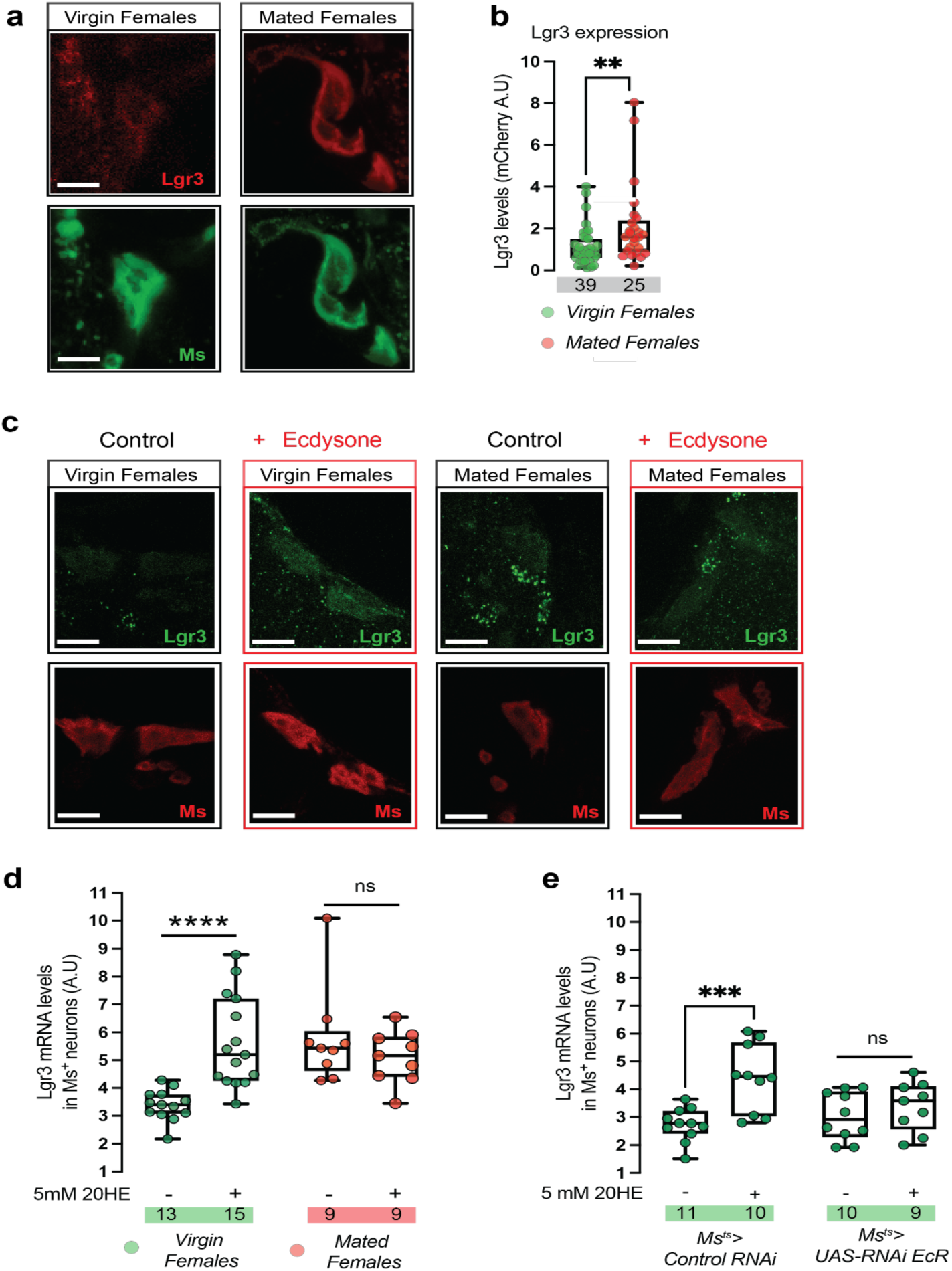
Relaxin receptor Lgr3 expression in Ms neurons is regulated by mating status and steroid hormone ecdysone signalling. **(a)** Representative confocal images showing *Lgr3* reporter expression (mCherry, red) and Ms neuron marker (Ms, green) in virgin females, and mated females. *Lgr3^T2A-Gal4^>UAS-mCherry* expression is increased in mated females compared to virgin females. Scale bars: 10 μm. **(b)** Quantification of Lgr3 (mCherry, red) in Ms neurons from (a), showing *Lgr3* levels. Mated females exhibit significantly higher mCherry intensity than virgin females (*p < 0.01). n values shown below. **(c)** In situ hybridization of endogenous *Lgr3* mRNA (green) in Ms neurons (red) from virgin females, and mated females, with or without 20HE treatment (red box outlines: +20HE). 20HE increases *Lgr3* expression in virgin females. Scale bars: 10 μm. **(d)** Quantification of *Lgr3* mRNA levels from panel (c). Treatment with 5 mM 20HE significantly increases *Lgr3* expression in virgin females (****p < 0.0001), but not in mated females (ns). n values are indicated in the boxplots. **(e)** Steroid signalling increases Lgr3 in Ms neurons via EcR. Quantification of Lgr3 mRNA in Ms⁺ enteric neurons after feeding 20HE to adult virgin females. Control flies (*Ms^ts^>Control RNAi*) show a significant ecdysone-induced increase in Lgr3 levels in Ms neurons (***p < 0.001). *Ms^ts^>UAS-EcR RNAi* flies significantly decreased the effect of ecdysone on Lgr3 levels.

Post-mating intestinal remodelling is coordinated by systemic endocrine signals, including the steroid hormone ecdysone^1,2^ and juvenile hormone (JH)^10^. We first asked whether JH signalling could prime Lgr3 expression in Ms neurons. We fed virgin females with the JH analogue methoprene (as previously described^10^) and measured *Lgr3* expression in Ms neurons using smRNA FISH. Methoprene treatment did not increase *Lgr3* mRNA levels in virgin females or mated females (**Extended Data Fig. 6c**). Thus, JH signalling alone is insufficient to promote Lgr3 expression in Ms enteric neurons.

We next examined ecdysone, the major steroid hormone in flies^1,2,10^, and a well-established orchestrator of post-mating physiological gut adaptations^1,11^. After mating in females, ecdysone levels rise systemically^11,36^ and trigger widespread intestinal remodelling by activating the nuclear ecdysone receptor (EcR) in both gut epithelial^11^ and neuronal tissues^1^. In particular, EcR signalling is required in Ms neurons for crop expandability, food intake and neuropeptide release^1^, placing ecdysone in a prime position to act upstream of Lgr3 expression.

To test whether ecdysone promotes Lgr3 expression in Ms neurons, we fed virgin and mated females with 20-hydroxyecdysone (20HE), as previously described^11^, and measured Lgr3 mRNA levels. Ecdysone feeding was sufficient to significantly increase Lgr3 expression in virgin females (**Fig. 3c,d**), while no effect was observed in mated females, suggesting that ecdysone can only boost Lgr3 expression before the mating-induced upregulation occurs. Consistent with an Ms-intrinsic, cell-autonomous mechanism, adult-restricted knockdown of EcR in Ms neurons abolished the ecdysone-induced increase in Lgr3 mRNA (**Fig. 3e**). Despite this increased Lgr3 expression, ecdysone-fed virgins did not exhibit crop enlargement (**Extended Data Fig. 6b**), indicating that ecdysone is not sufficient to trigger crop plasticity. Thus, ecdysone acts cell-autonomously in Ms neurons to upregulate Lgr3 expression after mating, preparing Ms neurons to respond to an additional mating-dependent signal that activates them to drive gut plasticity.

### dILP8 is a post-mating ovarian signal that activates enteric neurons via Lgr3

Having shown that Lgr3 is required in Ms neurons for reproductive gut plasticity, and that its expression is primed by ecdysone after mating, we next asked what ligand activates Lgr3 to trigger neuropeptide release and intestinal adaptations. Lgr3 is the known receptor for the relaxin-like ligand dILP8 in *Drosophila*^37,38^. In larvae, dILP8 activates defined Lgr3-positive brain neurons to delay metamorphosis when peripheral tissues are stressed^33,39^. We therefore tested whether dILP8 signal engages Lgr3 in adult enteric neurons to permit maternal intestinal plasticity. In adults, dILP8 has been implicated in control of ovulation^40,41^ and systemic metabolism^42^, but whether it regulates enteric neurons or gut plasticity remains unknown.

To determine if dILP8 could serve as a mating-dependent cue, we first examined its expression in virgin and mated females. Analysis of a previously published single-cell RNA-seq dataset from adult ovaries^43^ revealed high dILP8 expression specifically in terminal (stage 14) follicle cells surrounding fully mature oocytes (**Fig. 4a**). These cells are known sources of circulating reproductive hormones^44–49^. By using an *enhanced Green Fluorescent Protein* (*eGFP*) expressed along the first intron of dILP8, *Mi{MIC}dILP8^MI^*^00727^ (hereafter referred to as *dILP8-GFP)*^37,50,51^, we detected dILP8-GFP expression exclusively in late-stage (stage 14) follicle cells of the ovary (**Fig. 4b, Extended Data Fig. 7a-d**). These data place dILP8-expressing cells in a prime location to be relaying reproductive status to the GI tract.

**Figure 4.**
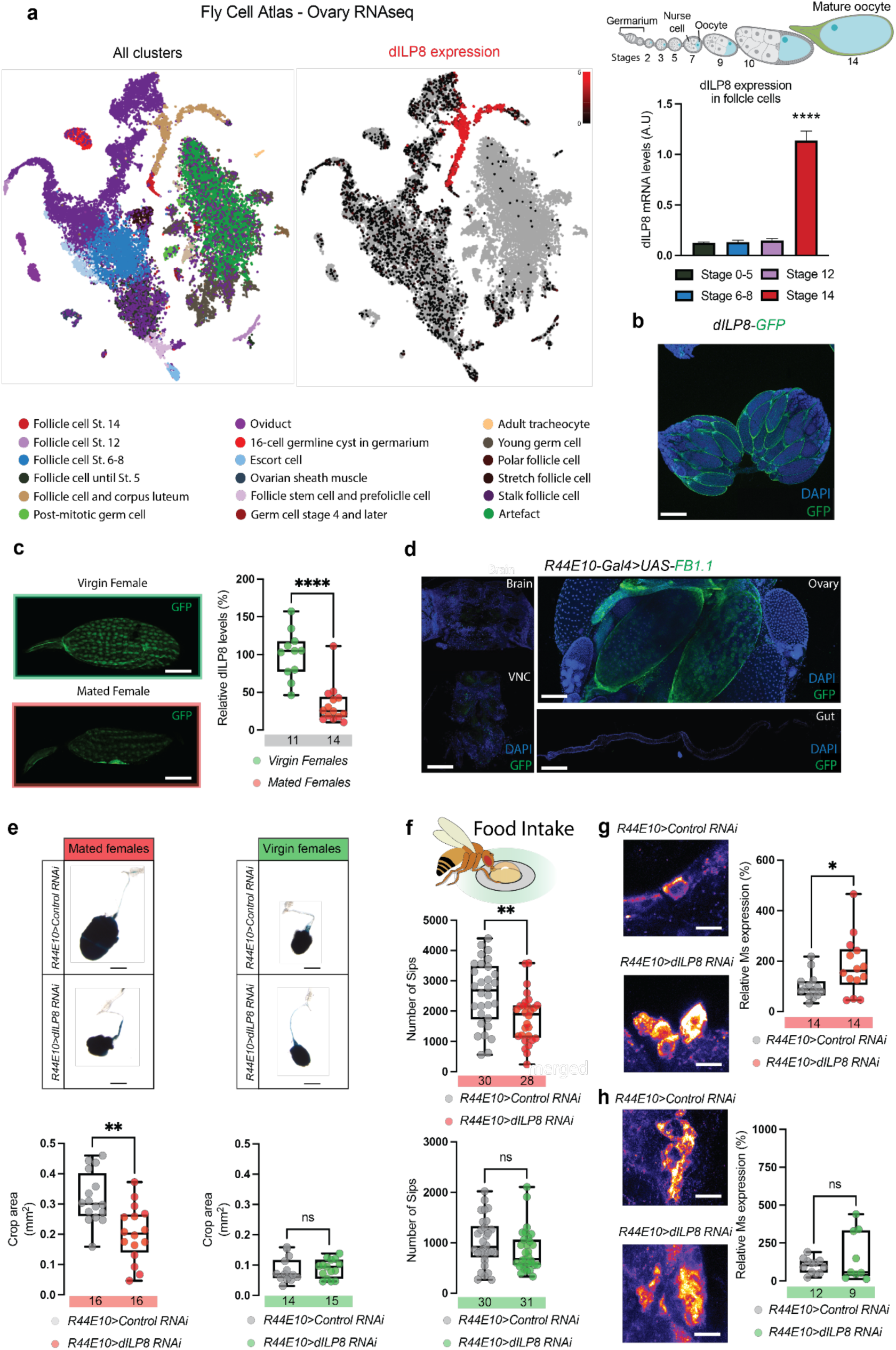
Ovarian dILP8 regulates female-specific crop plasticity and feeding behaviour via inter-organ signalling. **(a)** Left: UMAP plot from the Fly Cell Atlas RNA-seq^43^ of ovary cell types, highlighting all clusters. Middle: dILP8 expression is enriched in late-stage follicle cells. Right: Quantification of *dILP8* mRNA levels across follicle cell stages shows strong upregulation at stage 14 (****p < 0.0001). Top right: schematic of Drosophila egg development, from the germarium (where egg cells begin) to stage 14, the fully mature egg. **(b)** Confocal image of an ovary expressing *dILP8-GFP*, showing GFP signal in stage 14 follicle cells surrounding mature oocytes. Scale bar: 50 μm. **(c)** Confocal images and quantification of *dILP8-GFP* expression in stage 14 follicle cells from virgin vs. mated females. *dILP8* levels are significantly reduced in mated females (****p < 0.0001). Scale bar: 100 μm. **(d)** Tissue-specific expression pattern of the ovarian follicle cell driver *R44E10-Gal4>UAS-FB1.1*, showing strong GFP signal in the ovary follicle cells and no detectable expression in the brain, VNC, crop or midgut. Scale bars: 100 μm. **(e)** Representative crop images and quantification of crop size in mated females (red) and virgin females (green) upon *dILP8* knockdown in the ovary using *R44E10-Gal4*. Only mated females show a significant reduction in crop size (**p < 0.01); no effect is observed in virgin females. Scale bars: 200 μm. **(f)** Food intake quantified via FlyPAD. *dILP8* knockdown significantly reduces the number of sips in mated females (**p < 0.01), but has no effect in virgin females. **(g–h)** Confocal imaging of Ms peptide levels in Ms neurons of mated females (g) and virgin females (h) following ovary-specific *dILP8* knockdown. *Ms* expression is significantly upregulated in mated females (*p < 0.05) but unaffected in virgins (ns). Scale bars: 20 μm.

If dILP8 acts as a post-mating endocrine signal, its follicle-cell levels should change with mating. We therefore quantified dILP8 signal in ovaries from virgin and mated females. We found that dILP8 fluorescence was significantly reduced in mated females compared to virgins (**Fig. 4c**), indicating mating-status–sensitive expression, consistent with secretion upon oocyte maturation.

We next tested whether dILP8 is necessary for post-mating gut plasticity by downregulating *dILP8* expression specifically in mature follicle cells using the *R44E10-Gal4*^49^ driver (**Fig. 4d**). Targeted knockdown of dILP8 specifically in mature follicle cells impaired crop expandability in mated females (**Fig. 4e, Extended Data Fig. 8a**), but had no effect in virgin females. Consistently, post-mating increases in food intake were also reduced (**Fig. 4f**), indicating that dILP8 is required for the physiological adaptations triggered by mating.

Finally, we asked whether dILP8 acts through Lgr3 in Ms neurons. If so, dILP8 loss should prevent the neuronal activation required for Ms peptide release. Consistent with this, follicle cell-specific knockdown of dILP8 caused a strong accumulation of Ms peptide in Ms neurons of mated females (**Fig. 4g, Extended Data Fig. 8b**), but not in virgin females (**Figure 4h, Extended Data Fig. 8c**), phenocopying the effect of Lgr3 knockdown and consistent with dILP8 acting via Lgr3 to drive Ms peptide release.

Together, these data support a stepwise model: female genetic identity (*tra*) establishes Lgr3 competence in Ms neurons; after mating, the steroid hormone ecdysone primes these neurons by upregulating Lgr3 expression; and dILP8 released from stage 14 follicle cells activates Lgr3 to initiate maternal intestinal plasticity and increased food intake (**Fig. 5**).

**Figure 5.**
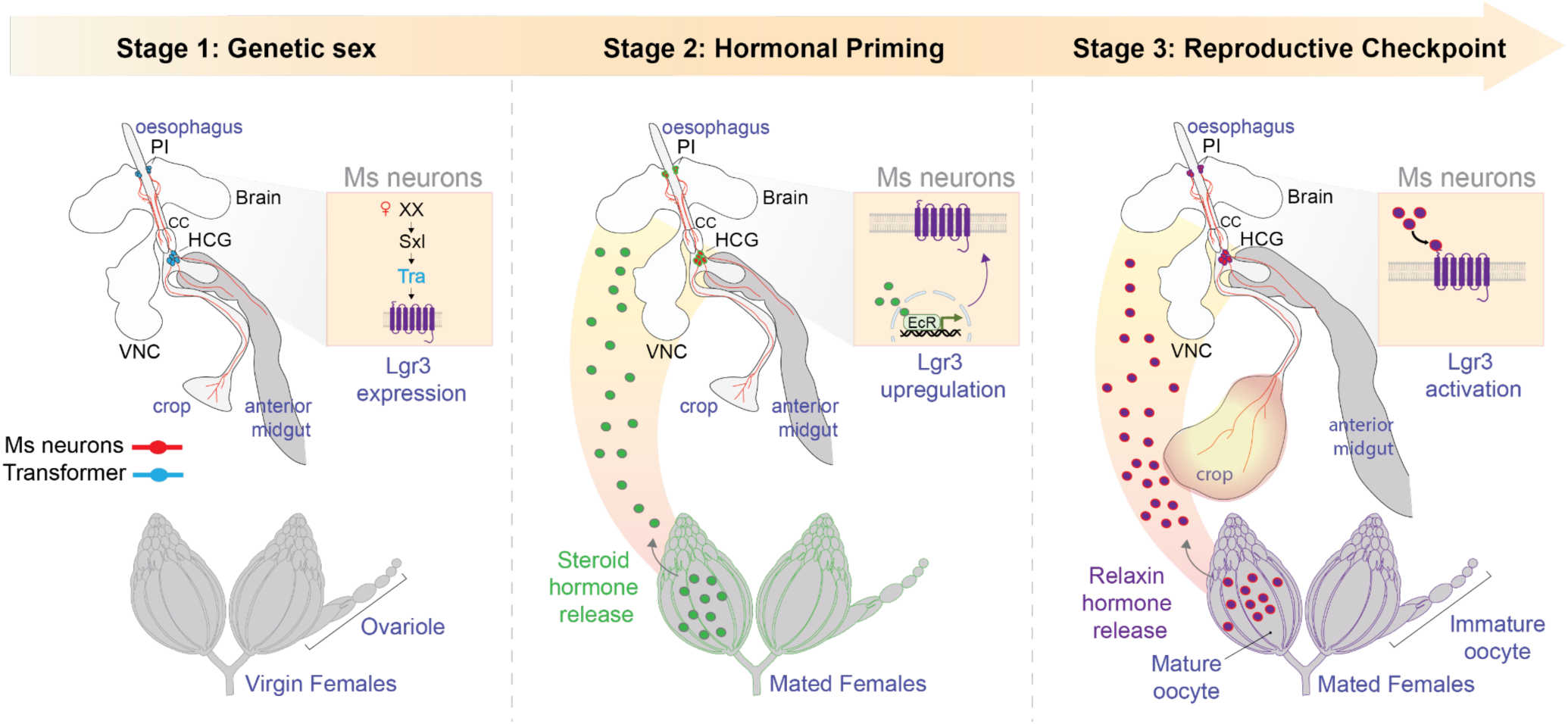
A three-stage neuroendocrine model linking reproductive state to gut plasticity via Ms enteric neurons in *Drosophila* females. Stage 1 – Genetic sex: In virgin females, the sex determination pathway, via *transformer (tra),* maintains female identity in Ms neurons (red), allowing expression of the relaxin-family receptor *Lgr3*. Stage 2 – Hormonal priming: After mating, ovarian ecdysteroids signal via ecdysone receptor (EcR)^1^ upregulate Lgr3, priming these enteric neurons for the reproductive signal while remaining in a primed, inactive state. Stage 3 – Reproductive checkpoint activation: Only after mating and once oocytes mature fully, follicle cells release the relaxin-like hormone dILP8, which activates Lgr3 on Ms neurons to switch on the reproductive checkpoint, coordinating intestinal plasticity and food intake with reproductive demand.

## Discussion

Successful reproduction requires coordination between energy balance and reproductive status. Here, we identify a neuroendocrine axis that gates the onset of intestinal adaptations in *Drosophila melanogaster*. The ovarian hormone dILP8 signals through its relaxin receptor Lgr3 in a subset of enteric neurons to induce gut enlargement and food intake. This circuit is female-specific and activated only after mating and oocyte maturation, defining a maternal intestinal plasticity checkpoint that permits these adaptations only once reproduction is underway (**Fig. 5**). Disrupting any step, sex determination, steroid priming or ovarian dILP8, prevents initiation despite mating.

Sex-specific physiology often requires sex-specific circuitry^52–55^. We find that the sex determination gene *tra* feminises enteric Ms neurons, endowing them with competence to express *Lgr3* and respond to ovarian cues. This female identity is specified developmentally and actively maintained in adulthood by *tra*, enabling female Ms neurons to decode systemic reproductive signals. Such persistent sexual dimorphism echoes principles observed in the central nervous system (CNS), where developmental sex determination cascades wire neural circuits for sex-specific behaviours^56,57^.

While Lgr3 was previously associated with developmental timing^31,33,38,39,58^, its role as a functional relaxin hormone receptor has not been demonstrated before. Here we uncover a novel role in adult visceral physiology. Ms neurons expressing Lgr3 are essential for reproductive intestinal enlargement and increased food intake, functions mediated through Ms neuropeptide release. However, Lgr3 activation is conditional. First, ecdysone, a steroid hormone secreted after mating, primes Ms neurons by upregulating Lgr3. Then, after mating and only upon full oocyte maturation, follicle cells secrete dILP8, which activates Lgr3. This hormonal relay ensures gut enlargement and increased food intake occur only when the female is reproductively committed. From an evolutionary perspective, this prevents unnecessary energetic investment and maximises fecundity.

After mating, ecdysone is secreted by immature follicle cells, triggering intestinal stem cell (ISC) divisions in the midgut via the ecdysone receptor (EcR/USP) signalling pathway^11^. In Ms neurons, EcR-dependent transcription likewise mediates the priming step (induction of Lgr3) thus establishing a “priming–gating” sequence. Whether an analogous steroid–peptide sequence gates ENS-mediated intestinal plasticity in vertebrates is unknown, although in the CNS oestrogen priming enables oxytocin-evoked maternal behaviour in rats^59^.

In mammals, relaxin-family peptides regulate uterine relaxation^60^, vascular remodelling^61^, and gut motility^62^. Our work demonstrates that in flies, dILP8 acts as a reproductive relaxin, coordinating visceral and metabolic changes with reproductive state. The shift in dILP8 function, from a local ovarian factor required for ovulation in virgins^40,63^ to a neuroendocrine signal after mating, suggests that dILP8 acts as a physiological checkpoint, gating maternal adaptations once reproduction is underway. Our model offers a point of comparison for the coordinated control of maternal metabolism in pregnant mammals^64^. In mice and humans, pregnancy hormones like progesterone, oestrogen, and relaxin not only remodel reproductive tissues but also reconfigure gastrointestinal physiology^62^, appetite^65^, and vascular tone^66^. Disruptions in these hormonal signals underlie maternal metabolic syndromes such as polycystic ovarian syndrome (PCOS)^67^ and pregnancy-associated insulin resistance^68^, emphasising the clinical relevance of decoding reproductive-metabolic crosstalk. Whether dILP8 and relaxin-family peptides act through analogous circuits, linking reproductive organs to enteric neurons, remains unknown. Nevertheless, similarities in timing (e.g., early and late pregnancy)^69^, function^62,70^, and receptor class^31^ strongly suggest conserved principles across phyla.

## Materials and methods

### Animals

Flies were raised on a standard cornmeal/agar diet (6.65% cornmeal, 7.15% dextrose, 5% yeast, 0.66% agar, supplemented with 2.2% nipagin and 3.4mL/L propionic acid). All flies have been outcrossed for at least five generations. All experimental flies were kept in incubators at 65% humidity and on a 12h light/dark cycle, at 18°C, 25°C or 29°C depending on the specific experiment. Flies were transferred to fresh vials every 2 days, and fly density was kept to a maximum of 20 flies per vial.

For transgene expression in Ms neurons, we used the *Ms-Gal4* line^71^. For transgene expression in stage 14 follicle cells, we used the *R44E10-Gal4* line (gift from Jianjun Sun)^49^. To restrict UAS/GAL4-mediated expression to the adult stage, we used the TARGET system^27^ with the *Tubulin-Gal80ts* line as previously described^72^. The following inducible driver lines were constructed in the laboratory and previously described: *Ms-Gal4;Tub-Gal80ts* (*Ms^ts^*)^1^; *UAS-GCaMP6f;Ms-Gal4*^1^. Gal4 activity was released by transferring 7-day-old adult flies to 29 °C for 3 d.

The following UAS-transgene lines were obtained from the Vienna Drosophila Resource Center (VDRC): *GD control* line (VDRC, 60000), *KK control* line (VDRC, 60100), *UAS-Tra RNAi GD764 (RNAi 1)* (VDRC, 2560), *UAS-dILP8 RNAi KK112161* (VDRC, v102604), *UAS-EcR RNAi GD1428* (VDRC, 37058). The following UAS-transgene lines were obtained from the Bloomington Drosophila Stock Center (BDSC): *Tubulin-Gal80ts* (BDSC, 7018), *attP40 control* line (BDSC, 36304), *attP2 control* line (BDSC, 36303), *UAS-Tra RNAi JF03132 (RNAi 2)* (BDSC, 28512), *UAS-Lgr3 RNAi GL01056 (RNAi 1)* (BDSC, 36887), *UAS-Lgr3 RNAi JF03217 (RNAi 2)* (BDSC, 28789), *UAS-Stinger* (BDSC, 84277), *UAS-SrcGFP* (BDSC, 5432), *UAS-mCherry.CAAX.S* (BDSC, 59021), and *UAS-GCaMP6f* (BDSC, 42747). Reporter lines used in this study include *Mi{MIC}dilp8^MI^*^00727^ (BDSC, 33079)^37,50^, *sfGFP::Lgr3 (ag5)*^33^, *TI{CRIMIC.TG4.2}Lgr3^CR00347-TG4.2^* (BDSC, 78895), *TraF^KI^-Gal4*^24^, and *UAS-Flybow*^1^. The *pHluorin-Ms* endogenous construct was generated in this study (see below for details).

4-day-old and 7-day-old virgin flies were used for experiments at 25°C and 18°C, respectively. Virgin female flies were either housed with males for 24h (mated) or kept as virgins for the control groups. In all experiments, mated female flies were dissected and analysed alongside age- and condition-matched virgin controls. For RNAi experiments, flies were reared and aged at permissive temperature (18°C) and then transferred to 29°C for RNAi induction for 3 days.

Mated female flies were mated 2 days following RNAi induction and tested on the third day of induction.

For the experiments using rodent primary neurons, we obtained brain samples from wild-type rats of the Sprague-Dawley strain Crl:CD(SD), bred globally by Charles River Laboratories in accordance with the International Genetic Standard Protocol (IGS). The execution of all experimental procedures strictly adhered to the guidelines set forth by the European Directive 2010/63/EU and the French Decree No. 2013-118, which concern the protection of animals used in scientific research. Approval for the experiments was obtained from the local ethics committee, Comité d’éthique N°5 Darwin.

#### Single-cell RNA sequencing

Single-cell transcriptomic data from dissociated fly midbrains, comprising eight biological replicates were obtained from^30^ and processed using Seurat version 5.2.1^73^. Male and female cells were distinguished on the expression of the long-non-coding RNA roX1 (expression > 0). To investigate sex-specific differences, cells with Ms expression above a log2-normalised value of 4 were selected for further analysis. Within this Ms-positive subset, cells were grouped by sex and biological replicate, and normalised counts were summed to generate pseudobulk profiles per replicate. Differential gene expression (DGE) between female and male Ms neurons was assessed using Seurat’s FindAllMarkers function (Wilcoxon rank–sum test on normalised expression). The dotplots in Fig. 2a show average expression (colour) and abundance (size) across Ms-positive neurons in males and females for the 12 genes most significantly upregulated in females, ranked by adjusted p-value from the Seurat FindAllMarkers results.

#### Immunohistochemistry

Adult Drosophila brains and guts were dissected in PBS and mounted on poly-L-lysine-coated slides (Sigma-Aldrich, P1524). Tissues were fixed at room temperature in 4% paraformaldehyde (PFA) in PBS for 40 minutes. After fixation, tissues were rinsed briefly in PBS to remove residual fixative. Then, samples were permeabilized in PBS containing 0.2% Triton X-100 (PBT) for 15 minutes and then blocked in PBS with 4% horse serum and 0.2% Triton X-100 (PBTN) for 1 hour at room temperature. Primary antibodies were applied in PBTN and incubated overnight at 4°C. The following primary antibodies were used: goat anti-GFP (1:1000, Abcam, ab5450), mouse anti-RFP (1:200, MA5-15257, ThermoFisher Scientific), rabbit anti-Ms (1:1000, K7834, New England Peptide) and Alexa Fluor 647 Phalloidin (1:300, ThermoFisher Scientific, A22287).

Following primary antibody incubation, samples were washed in PBT (3-4 washes, 5 minutes each) at room temperature. We used the following secondary antibodies: Alexa Fluor 488 goat anti-mouse (A11029, Invitrogen), Alexa Fluor 488 goat anti-rabbit (A11034, Invitrogen), Alexa Fluor 596 goat anti-mouse (A11005, Invitrogen), Alexa Fluor 596 goat anti-rabbit (A11037, Invitrogen), Alexa Fluor 488 donkey anti-goat (A11055, Invitrogen) and Alexa Fluor 633 goat anti-mouse (A21126, Invitrogen). Secondary antibodies were then applied in PBTN and incubated for 1.5 hours at room temperature. Samples were washed in PBT (3 washes, 5 minutes each) and then three times in PBS. Finally, tissues were mounted in Vectashield (Vector Laboratories, H1200) and visualised under a Stellaris Confocal microscope.

#### RNA Fluorescence In Situ Hybridization

We have used the RNAscope Multiplex Fluorescence Detection Reagents v2 (Advanced Cell Diagnostics, 323110) and RNAscope H2O2 and Protease Reagents, Advanced Cell Diagnostics, 322381). All surfaces and equipment were cleaned with RNAzap (Invitrogen, AM9780) prior to dissection. Brains were dissected in RNAlater (Sigma-Aldrich, R0901) diluted 1:50 in DEPC-treated water and briefly rinsed in 1X PBS (prepared in ultrapure water) (ThermoFisher Scientific, 70011044). Dissected tissues were mounted on poly-L-lysine-coated slides in 1X PBS and fixed immediately in cold 4% formaldehyde (FA) (Sigma-Aldrich, F8775) for 30 minutes. Following fixation, FA was removed, and tissues were incubated with 2–3 drops of hydrogen peroxide for 10 minutes at room temperature (RT) until foaming was observed. Tissues were then washed three times for 5 minutes each in wash buffer. Wash buffer (Advanced Cell Diagnostics, 320058) was prepared by diluting 1:50 in ultrapure water and preheated to 40°C for 10 minutes.

##### Dehydration

Slides were sequentially incubated with 200 μL of ethanol solutions as follows: 50% ethanol for 5 minutes at RT, 70% ethanol for 5 minutes at RT and 100% ethanol for 5 minutes at RT (repeated twice). After the final ethanol step, slides were air-dried for 5 minutes before applying 5 drops of Protease III per slide, followed by a 30-minute incubation at RT.

##### Probe Hybridization

RNAscope FISH probes were preheated to 40°C for 10 minutes and cooled at RT for 5 minutes before use. Probes were prepared by combining 50 μL of C1 (Dm-Lgr3, Advanced Cell Diagnostics, 424001) with 1 μL of C2 (Dm-Ms-C2, Advanced Cell Diagnostics, 516011-C2). Tissues were washed three times in 1X PBS (1 mL per wash, 5 minutes each). During the second wash, probes were placed in the oven at 40°C, and during the third wash, probes were allowed to cool to RT. After removing PBS, FISH probes were applied to tissues, and slides were placed in a humidified chamber for hybridization at 40°C for 2 hours.

##### Signal Amplification

Slides were subjected to hybridization with AMP reagents in sequential steps. AMP1: Warmed to RT, applied to slides (4 drops/slide), and incubated in the oven at 40°C for 30 minutes. Tissues were washed three times in wash buffer for 5 minutes each. Then, AMP2: Warmed to RT, applied to slides (4 drops/slide), and incubated at 40°C for 30 minutes. Washes were repeated as described. Next, AMP3: Warmed to RT, applied to slides (4 drops/slide), and incubated at 40°C for 15 minutes. Washes were repeated as described.

##### Detection with HRP-Conjugated Probes and Opal Dyes

HRP-conjugated probes were applied depending on the channel to be revealed (C1, C2). For each channel, slides were incubated at 40°C for 15 minutes. After washing three times in wash buffer, Opal dyes (1:1000 in TSA multiplex buffer, Advanced Cell Diagnostics, 322809) were applied to slides and incubated at 40°C for 30 minutes. The Opal dyes used were as follows: 520 nm (Akoya Biosciences, OP-001001) and 570 nm (Akoya Biosciences, OP-001003). Following incubation, slides were washed and treated with HRP blocker (4 drops/slide) for 15 minutes at 40°C. Washes were repeated, and the process was repeated for each probe.

##### Counterstaining and Mounting

Tissues were counterstained with DAPI for 5 min at RT and washed once in the wash buffer. Slides were mounted using Vectashield mounting medium (Vector Laboratories, H1000).

#### Food intake assay

FlyPAD assays were performed as previously described^1,17,74^. In each FlyPAD arena, half of the electrode wells were filled with a pellet of cornmeal/agar diet, which were prepared by using a 1ml pipette tip to punch out pellets to match the diameter of the inner electrode circle. The remaining wells were left empty to monitor non-feeding baseline interactions. Flies were given 1h30 to feed in the arenas, maintained at 25 °C and 70% humidity. Capacitance was recorded using Bonsai software^74^, and the total number of sips per fly during the 1h30 feeding session was extracted using a custom MATLAB script^74^. All FlyPad experiments were carried out between 10:00 AM and 13:00 PM, and data from experimental and control groups were collected within the same assay to ensure consistency.

#### Crop size mounting and measurements

Crop size was measured in response to a standard food diet, as previously described^1^. Virgin female flies were collected and aged for 4 or 7 days, depending on whether they were raised at 25 °C or 18 °C, with regular transfers to fresh food every 2 or 3 days respectively. Following this period, some flies were mated for 24 hours, while others remained as virgin controls. After mating, all flies were given unrestricted access to standard food. The following morning at 10:00, flies were gently transferred to tubes containing FCF Blue food and allowed to feed freely for 1 hour and 30 minutes. They were then moved to an empty vial by tapping, euthanized using snap freezing in liquid nitrogen, and either dissected immediately or stored at −80 °C for later analysis. No tissues were thawed and re-frozen during this process. For each experiment, both experimental and control flies were raised and tested under the same food conditions. Tissues were dissected in 1× PBS and transferred to slides for immediate bright-field imaging. Crop area was evaluated on segmented crops using the Fiji image analysis software.

#### In vivo calcium imaging

In vivo imaging experiments were carried out as previously described^1,72^. To monitor Ca^2+^ activity levels, females flies carrying a *UAS-GCaMP6f;Ms-Gal4* were crossed to control males or to males carrying the appropriate *UAS-RNAi Tra GD764* or *UAS-RNAi Lgr3 GL01056* for imaging in Ms enteric neurons. Crosses for imaging experiments were raised at 22°C. Briefly, a single fly was affixed to a plastic coverslip through a dental glue (Protemp II 3 M ESPE). Then 100 μl of an artificial haemolymph solution was added on top of the coverslip. The composition of the artificial haemolymph solution was: 130 mM NaCl (Sigma, S9625), 5 mM KCl (Sigma, P3911), 2 mM MgCl2 (Sigma, M9272), 2 mM CaCl2 (Sigma, C3881), 5 mM d-trehalose (Sigma, T9531), 30 mM sucrose (Sigma, S9378) and 5 mM HEPES-hemisodium salt (Sigma, H7637). Surgery was performed as previously described^72,75^, to expose the brain for optical imaging. Then, a fresh drop of 100 μl of the haemolymph solution was applied on the top of the opened fly head cuticle.

Two-photon imaging was performed on a Leica Stellaris 8 DIVE upright microscope equipped with a ×25, 1.00-NA water-immersion objective. Two-photon excitation of GCaMP6f was achieved using a Mai Tai eHP DeepSee MP laser tuned to 920 nm. Detectors were tuned for GCaMP6f at 500-550nm. Then, 512 × 512 images were acquired at a rate of one image every 648 ms in a single plane, and the entire duration of each recording was 300s. Image analysis was performed offline with a graphical user interface, custom-programmed with MATLAB. Regions of interest (ROI) were delimited by hand and surrounding individual cell bodies of GCaMP6-expressing cells. Amplitude oscillation measurements were conducted as described^1,76^. Each dot represents the average of individual neuronal somas of a single fly brain.

#### pHluorin-Ms cloning

*pHluorin-Ms* construct flies were generated using CRISPR–Cas9-assisted homologous recombination as described^77^. The entire coding region of the gene was removed and replaced by pHluorin-Ms. For the homologous recombination two-guideRNAs inserted in the pCFD4 vector (Addgene, 49411) were used gRNA1: 5′-TTTTAGAGCTAGAAATAG-3′ and gRNA2: 5′-AACACCACTTGGTCCCGA-3′. pTV3-Ms vector, described in^1^, which contains the two homology arms 5’ and 3’ of the coding region for *Myosuppressin*, was digested with restriction enzymes NdeI and BfuAI. pHluorin was PCR amplified from the plasmid CaMKII-Ms-pHluorin using the primers forward 5’-cacacgggccGGATCTAAAGGCGAGGAAC-3’ and reverse 5’- cctgaactgcTTTGTACAGCTCGTCCATG-3’. The genomic locus of Ms was amplified from fly genomic DNA. The first intron and the ER signal of Ms were amplified using the primers forward 5’-ttaatcaaaatcagtggctaGGTTGGATGGACATGGGG-3’ and reverse 5’- ctttagatccGGCCCGTGTGTTGGACAC-3’ while the rest of the Ms genomic region was amplified using the primers forward 5’- gctgtacaaaGCAGTTCAGGGTCCACCTC-3’ and reverse 5’- ggggcactacggtacctgcaTTAACGACGTTTTCCGAAACGC-3’. All the fragments were assembled in the opened pTV3 plasmid using Gibson assembly^78^, #E5520 NEBuilder HiFi DNA Assembly Cloning Kit, to form the pTV3-ER Signal-pHluorin-Ms plasmid. pTV3 and pCFD4 plasmids were injected into yw; nos Cas9 (II-attP40) flies by BestGene.

5’ homology arm: gtgcttgcgttcaacaagtccagcaaacagagcagcagctgaaccccggtgttaacaactaacaagtttgtccattaacttctttgtggaagcaccgatacctcaaagccctcatcaggtgggtacttgtgtcttgagatgtgcagagtgatagatactttagaggaataactgaatacatataaagtgaatccttgaggtttcagtcgaaaggtgtgaaagataaagcctgtattaaaagtgtgtacatttgtgaaaatatggtactatcataatgatggctttatacttaattattcaattatccaacgaatatcaccagcttgcctggtcttgtaagaatgattagaaaatttggtattttggtatttaaaagaatggtagaattgcgctaatataaagtgtaaagctatttaaaaatagtccaaaaacgtaaggtagatgaagtggaaagtattgtagtttttaaaaacgctatggtatgtgaggaagatttcctataaatatgtaatttaacatttaaagaacttattagttttgaccatgagtgatagacatttcaactaagtcgcaatagatggtttcttgtgagtaaacagacatggcaattgatttgcatacgtgcaccttgattgagccctaaacaagcatcagtagtttggatccttggaacgtgtcctatgtgcaactcccgcccggcatctactcctccctccagacttccggtgctggtttttcta agctaaacagatgtgggaacaacacgttcgcacaggtgtttgcatgccgactgcaacacggggcgtatgagtgctgcctccacttccatcatttcgagcgtaatcatcatcccgaggcgttgacgcagaacaaattgccttagcctccgccattttcagctaatagaaacaaattgtgtgtcgcgtaaacgtattagggtaccattaagacgcctgcttggatgcgatttaaaatggtaacaccgccgctagccagaaggccaagtacaactccatttatgcataatactttgccagggcaacgccatcatcagcgaatggcaatcaggcacgtagcattaagatcattacacttaatcaaaatcagtgg

3’ homology arm: ccgacatgagacaacgacactggaccctgaccacaagcggcggaatcgtttctgttcacccaaaaagcacaacactattttgacgtcttcagcataattatgtaaacgtaatcgatggaaactcagaactatactcaattggaagctctctagttcattaaatatccaatgtccaatgtttctatgcaacaaaaaaaaaatcgaatacatatttgtaaatactcaaagaccctcgaaatgttctgaaagttaaacccttggttttgatttaattcgtactctttatttgctgagtgttataaagaactaataatacgtatttcaacgatgtttaaatatctcacacatatttccctagcatgaagcactattattaaataaccaacaaatgttttcaaatccaaacactattttccgttgtatactttaataaagacaaacttttcctctcaatttgtgaatgcatagcaaaatgcaattgaaatggtttacatttaataggaaagttgggctactctttgaacaacattcaacaacaatgattttggcgagttagattgtgaacttcatacataattaacttttttgctcctttctaacaagtttatatagtcaatcaccatggaataaacaataaaaaaaggtacgaaatttttttttacattttaatttattactgttgacggtttcttatacgttaaaacattctaataaagtcaattttactaaatggattattgacgctattgcattttgttgtacgtcatttgcgtaatctttgaaaaatatttccgaattttattcgtatccttgaaatataatttcgtatgagaatggttaatgggttccatagtacgcagatattttcgctccattgggttttttgattttcaatttttttgcttttgctgaaaaagttaaaagttaccatttaattgcatgtttttattaaattattttgccattcttaaaggttttatttaaattaataaaaaattaaacaaataacagaatattctaaatcaaatggacagaaaaacgtgaaataatgcagttattattcataaaatgtctagacttgcaaattaaaaattgtatgacttttaaaaattagtttctttgtctgattctcattacatattgcc

#### Ms-pHluorin validation in primary cortical neurons

Ms-pHluorin translocation assays *in vitro* were performed in primary co-cultures of cortical neurons and astrocytes by transfecting Ms-pHluorin sparsely in a few neurons using the calcium-phosphate method, as done previously^76,79^. Experiments were carried out from DIV12 to DIV21 and axonal regions were identified by the co-expression of Synapsin-mRuby3^29^. Imaging was performed on a custom laser-illuminated epifluorescence microscope (Zeiss Axio Observer 3) with an Andor iXon Life camera (IXON-L-897) cooled to -70°C. Fiber-coupled lasers at 488 nm (Coherent OBIS LX, 30 mW) and 561 nm (Coherent OBIS LS, 80 mW) were combined with a Coherent Galaxy beam combiner and gated by a custom Arduino-based circuit synchronizing illumination and imaging. Neuron-astrocyte co-cultures on coverslips were mounted in a closed-bath stimulation chamber (Warner RC-21BRFS) and imaged through a 40x Zeiss Plan-Neofluar oil objective (NA 1.30; WD 0.21 mm). Field stimulation used 1 ms current pulses between platinum-iridium electrodes in the chamber (Warner PH-2), driven by a stimulus isolator (WPI A382) under control of the same Arduino circuit. Fluorescence changes were acquired at 10 Hz. All experiments were performed at 36.5°C, maintained with a dual-channel temperature controller (Warner TC-344C) that heated the stimulation chamber (PH-2) and the perfusate via an in-line heater (Warner SHM-6). Solutions were perfused at 0.35–0.40 ml/min. Tyrodes contained (mM): 119 NaCl (Merck, 31434), 2.5 KCl (Sigma, P9333), 2 CaCl_2_ (Sigma, C1016), 2 MgCl_2_ (Sigma, 68475), 20 glucose (Sigma, G8270), and 25 HEPES (Sigma, H3375) (pH 7.4 at 37°C), with 10 uM CNQX (Sigma, C-141) and 50 uM AP5 (Sigma, D-145). Trafficking of Ms-pHluorin was induced by field stimulation, using 400 pulses at 100Hz. Analyzing ΔF/F0 of individual boutons showed dramatic changes in fluorescence compatible with single vesicle exocytosis.

#### In vivo pHluorin-Ms imaging

*In vivo pHluorin-Ms* experiments were carried out as previously described^1,29,72^. To monitor *pHluorin-Ms* activity levels, 4d old virgin or mated female flies carrying the endogenous *pHluorin-Ms* construct were used for imaging in Ms enteric neurons. Flies for imaging experiments were raised at 25°C. Imaging solution and surgery was performed similarly as for calcium imaging experiments. Imaging recordings were carried out in the same set up as for calcium imaging experiments. Two-photon excitation of pHluorin was achieved using a Mai Tai eHP DeepSee MP laser tuned to 920 nm. Then, 512 × 512 images were acquired at a rate of one image every 648 ms in a single plane, and the entire duration of each recording was 600s. Image analysis was performed offline with a graphical user interface, custom-programmed with MATLAB. Regions of interest (ROI) were delimited by hand and surrounding individual cell bodies of *pHluorin-Ms*-expressing cells. Amplitude measurements were conducted as previously described^1,76^.

#### dILP8 transcript levels

Relative expression of *dILP8* in follicle cells at different stages was obtained from published single-cell transcriptomic data^43^, according to clustering performed in the original study where follicle cells are categorised into stage 0-5, stage 6-8, stage 12 and stage 14.

#### Drug feeding assay

Ecdysone or Methoprene feeding experiments were conducted following a previously described protocol^11^. Briefly, virgin or mated females were used for these assays. Ecdysone (Sigma-Aldrich H5142) or Methoprene (Supelco 33375) were first dissolved in 100% ethanol, and then diluted with water to prepare a 25 mM stock solution in 10% ethanol. For feeding assays, a final concentration of 5 mM drug was used, while control flies received a solution containing 2% ethanol. A mixture containing 50 μl of 5% sucrose solution, 5 mg/ml dry yeast and 5 mM drug was applied onto a Whatman filter paper, place inside an empty vial. The control group received the same sucrose-yeast mixture but with 2% ethanol instead of drug.

#### Statistics

All statistical analyses were carried out in GraphPad Prism 10. Behavioural comparisons between two genotypes and/or conditions were analysed with the Mann-Whitney-Wilcoxon rank sum test, as previously described^17^. The Mann-Whitney-Wilcoxon rank sum test does not require the assumption of normal distributions, so no methods were used to determine whether the data met such assumptions. Confocal images were compared either via t-test or one-way ANOVA. Live imaging experiments were compared via t-test. All graphs were generated using GraphPad Prism 10.

All confocal and bright field images belonging to the same experiment and displayed together in our figures were acquired using the exact same settings. For visualization purposes, level, and channel adjustments were applied using ImageJ to the confocal images shown in the figure panels (the same correction was applied to all images belonging to the same experiment), but all quantitative analyses were carried out on unadjusted raw images.

In all figures, n denotes the number of brains, guts or group of flies that were analysed for each genotype. Data are presented as boxplots with all data points shown, p values from Mann-Whitney-Wilcoxon test, t-test or one-way ANOVA (non-significant (ns); *p<0.05; ** p<0.01; ***p<0.001, ****p<0.0001). Asterisks highlighting significant comparisons across sexes are displayed in grey boxes, whereas those highlighting significant comparisons within same-sex datasets are displayed in red (for mated females) and green (for virgin females).

## Acknowledgements

We thank Pierre Leopold, Alisson Gontijo, Jianjun Sun, the Bloomington *Drosophila* Stock Center (BDSC) and the Vienna *Drosophila* Resource Center (VDRC) for fly stocks. We thank the LMS Microscopy and the Crick Advanced Light Microscopy STP for support. Part of this work was carried out in the ICM.Quant core facility of ICM, we gratefully acknowledge Claire Lovo and Astou Tangara for support. We are grateful to Alexandra Milona and Alexandros Kontopoulos for helping us with reagents and lab management. We thank Alessandro Mineo and Chris Amourda for helpful daily discussions throughout this research. We thank Claire Wyart for sharing antibodies with us. We are grateful to members of the Organ Development and Physiology Laboratory and GutSense laboratories for comments on the manuscript. We are grateful to Venizelos Papayannopoulos, Bassem Hassan, Nelson Rebola, Nikolaus Konstantinides, Katarzyna Siudeja, Clement Carre, Michael Rera, Celine Cansell and Zhou Xu for comments on the manuscript.

## Funding

ERC Starting Grant ( ERCStG 101117267 ‘GutSense;, DH)

Paris Brain Institute (ICM) intramural funding (DH). Paris Brain Institute Diane Barriere Chair in Synaptic Bioenergetics awarded to J.d.J.-S., funding from the French national program “Investissements d’avenir” ANR-10-IAIHU-0006 awarded to the ICM and funding by the Richard Mille Fund (Project DBS, 2023-2028).

Fondecyt Grant No 1231556 (JMC)

ERC Starting Grant (ERC-StG-852873 “SynaptoEnergy, J.d.J.-S.)

Big Brain Theory (BBT4, DH)

ERC Advanced Grant (ERCAdG 787470 ‘IntraGutSex’, IM-A), UKRI Frontier Research Guarantee grant (EP/Y036298/1, IM-A)

MRC intramural funding (IM-A)

S.W was financed by Wellcome Principal Research Fellowship (200846), a Wellcome Discovery Award (225192), an ERC Advanced Grant (789274), and Wellcome Collaborative Awards (203261 and 209235).

UKRI Horizon Europe funding Guarantee [EP/X038882/1 ’InnovaTEbehaviour’, CDT] Francis Crick Institute, which receives its core funding from Cancer Research UK (CC2258), the UK Medical Research Council (CC2258) and the Wellcome Trust (CC2258UK-China Research and Innovation Partnership Fund through the Met Office Climate Science for Service Partnership (CSSP) China as part of the Newton Fund, IM-A

## Author contributions

Conceptualization: BS, DH, IM-A

Methodology: BS, ET, JJ-S, DH, IM-A

Investigation: BS, ET, PG, AK, JD, AH, CR, JJ-S, JMC, CT, SW, DH

Visualization: BS

Funding acquisition: DH, IM-A

Project administration: BS, DH, IM-A

Supervision: DH, IM-A

Writing – original draft: BS

Review and editing: BS, ET, AK, JD, PG, AH, CR, JJ-S, JMC, CT, SW, DH, IM-A.

## Data availability

All other data, new material are available by the corresponding author upon reasonable request, or are present in supplementary data files.

## Extended Figures

**Extended Data Fig. 1.**
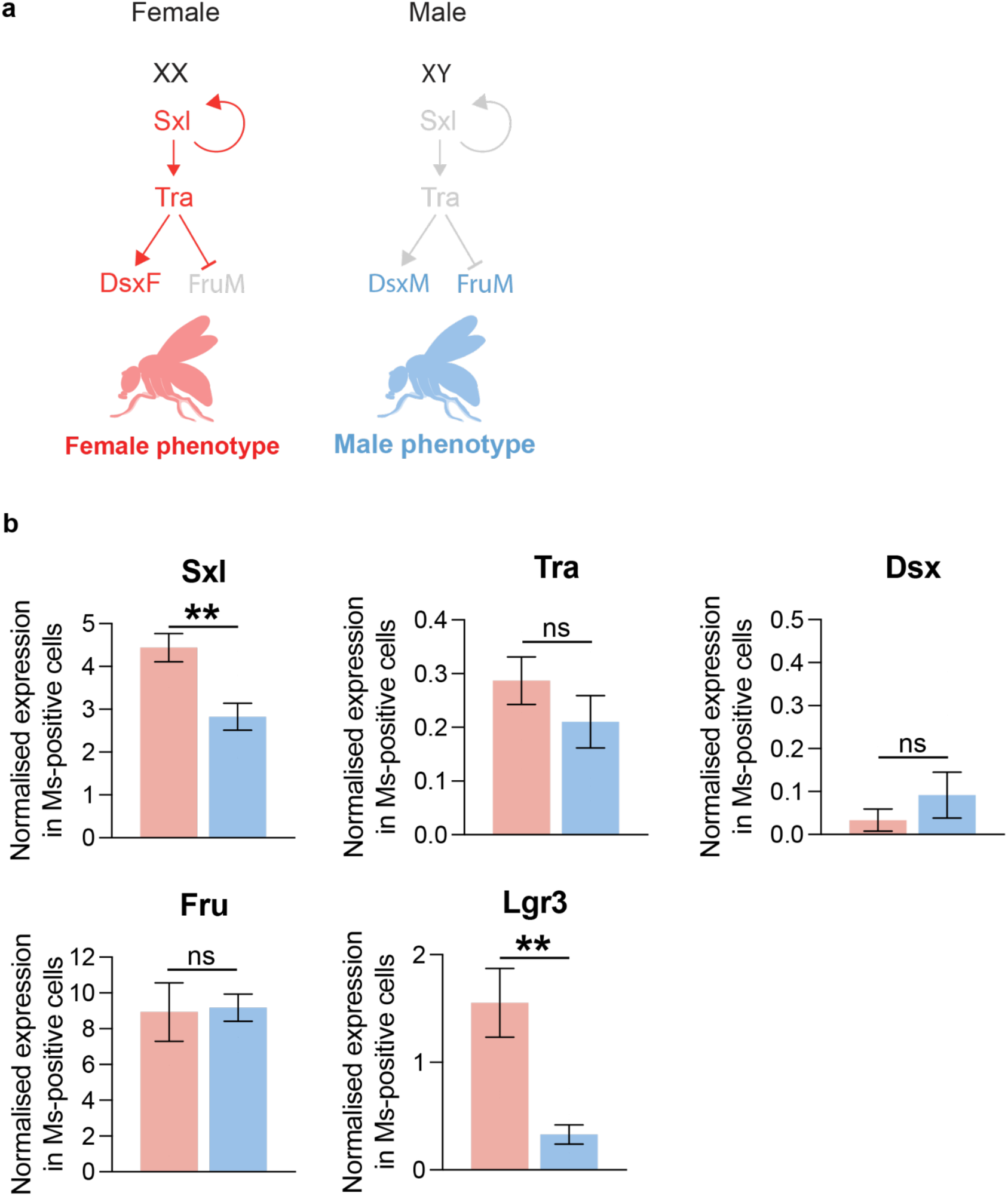
Canonical sex determination pathway and sex-specific expression of pathway components in adult Ms neurons. **(a)** Schematic of the Drosophila sex determination pathway cascade. In XX females, activation of Sex-lethal (Sxl) and its positive feedback loop maintains Sxl expression and promotes female-specific splicing of *transformer* (*tra*), generating functional Tra protein. Tra then directs female-specific splicing of *doublesex* (*dsx*) into the female isoform DsxF and prevents production of the male-specific FruM isoform from *fruitless* (*fru*), resulting in a female sexual phenotype. By contrast, in XY males, Sxl is not activated, tra transcripts are not spliced, and default splicing produces DsxM and FruM, leading to a male sexual phenotype. **(b)** Normalised pseudobulk expression of *Sxl, tra*, *dsx*, *fru* and *Lgr3* in adult Ms neurons from female (red) and male (blue) brains. Pseudobulk values were obtained by summing expressions across all Ms-positive cells per sex and normalising for library size. Among the candidates tested, *Lgr3* and *Sxl* show clear female-biased expression (**p < 0.01), whereas *tra* and *dsx* are barely detectable (ns) and *fru* is expressed at similar levels in both sexes (ns). Bars show mean normalised expression; error bars indicate s.e.m.

**Extended Data Fig. 2.**
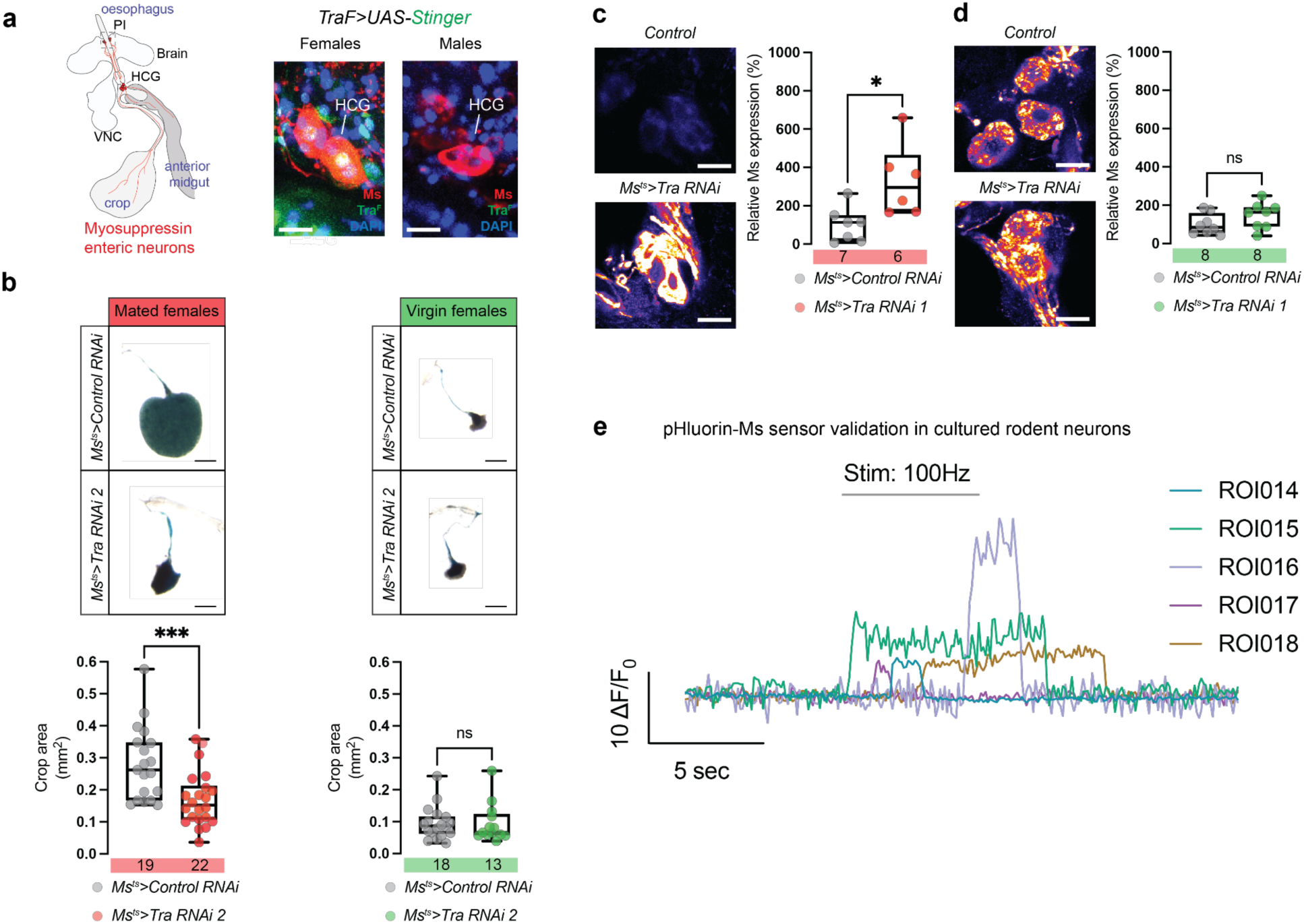
Independent *Tra* RNAi line confirms female-specific effects on crop size and enteric Ms expression. **(a)** Left: Diagram of Ms neuron projections along the gut. Right: Confocal images showing nuclear GFP (green) in Ms neurons (red) of the hypocerebral ganglion (HCG) in *TraF^KI^>UAS-Stinger* flies. GFP signal is observed in females but absent in males, confirming female-specific expression. Scale bars: 10 μm. **(b)** Representative crop images (top) and quantification of crop area (bottom) in mated females (red) and virgin females (green) expressing a second *Tra* RNAi line (*Tra RNAi 2*) under *Ms*-Gal4. Knockdown reduces crop size specifically in mated females (***p < 0.001), with no effect in virgin females (ns). Scale bars: 200 μm. **(c–d)** Confocal images for *Ms* peptide levels in the HCG of mated (C) and virgin (D) females. Ms peptide expression is higher in mated females upon *Tra* knockdown (*p < 0.05), but remains unchanged in virgin females (ns). Scale bars: 10 μm. **(e)** pHluorin-Ms sensor validation in cultured rodent neurons. Representative ΔF/F₀ traces from individual ROIs during field stimulation (100 Hz; grey bar). Stimulation evokes transient increases in pHluorin-Ms fluorescence, consistent with exocytosis-dependent dequenching. Colours denote distinct ROIs. Scale bars: 10% ΔF/F₀, 5 s.

**Extended Data Fig. 3.**
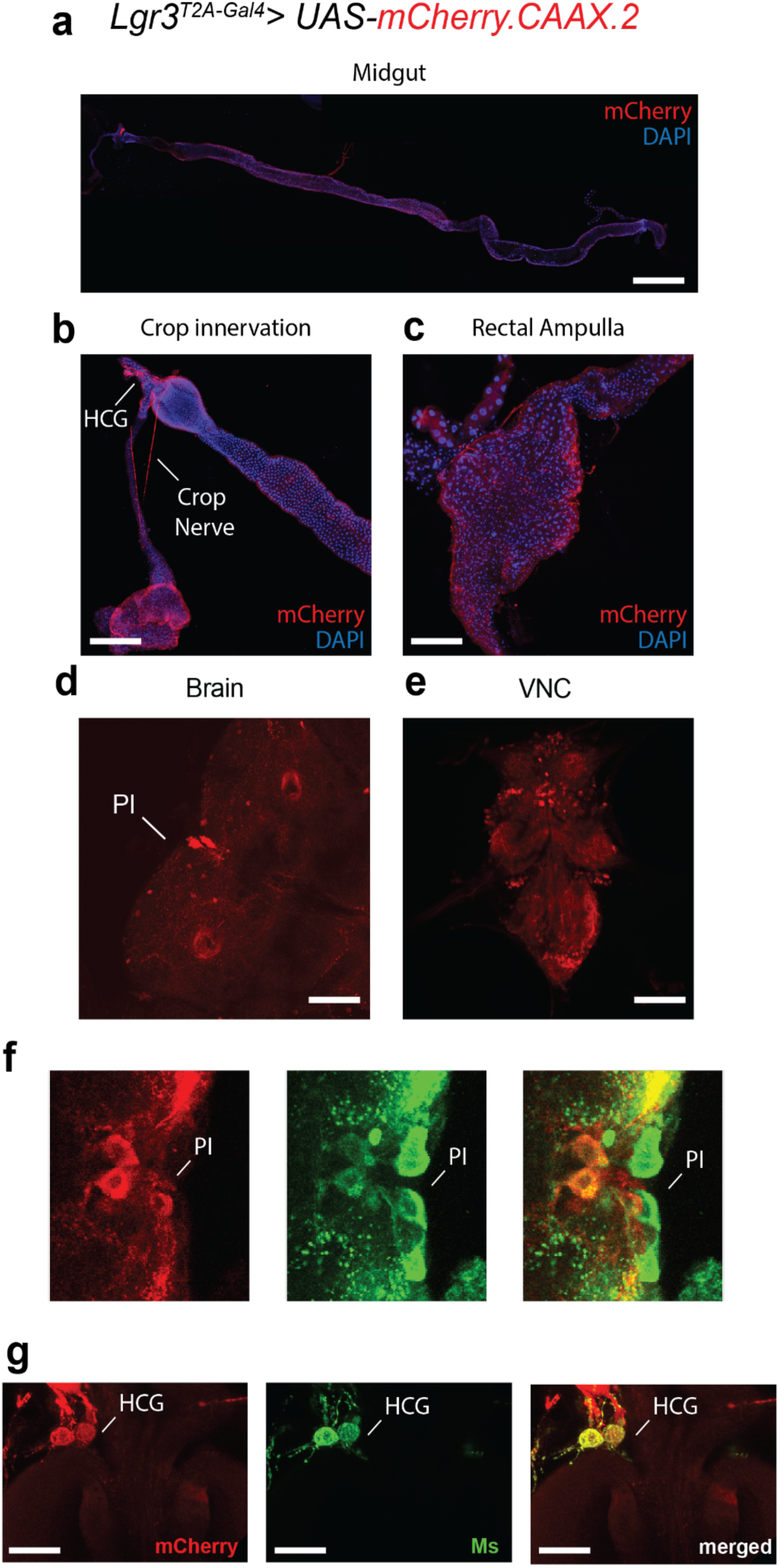
Expression pattern of *Lgr3* in the adult female *Drosophila* nervous system and gut. **(a)** Whole midgut preparation showing *Lgr3*-driven membrane mCherry signal (red) along the anterior-posterior axis. Nuclei are labeled with DAPI (blue). Scale bar: 200 μm. **(b)** Crop region showing innervation of the crop nerve and hypocerebral ganglion (HCG) by *Lgr3*-positive fibers. mCherry marks cell membranes. Scale bar: 50 μm. **(c)** Rectal ampulla with dense *Lgr3*-driven mCherry expression. Scale bar: 50 μm. **(d–e)** Brain and ventral nerve cord (VNC) expression patterns. In the brain (d), mCherry expression is visible in the pars intercerebralis (PI); in the VNC (e), scattered *Lgr3*-positive cells are observed. Scale bars: 50 μm. **(f)** Co-labeling of mCherry (red) and Ms neurons (green) in the PI region confirms overlap of *Lgr3*-expressing neurons with Ms neurons. Right panel shows merged image. Scale bar: 20 μm. **(g)** Co-localisation of *Lgr3*-driven mCherry (red) and Ms neuron marker (green) in the HCG. Merge indicates Lgr3 expression in Ms neurons within the HCG. Scale bar: 20 μm.

**Extended Data Fig. 4.**
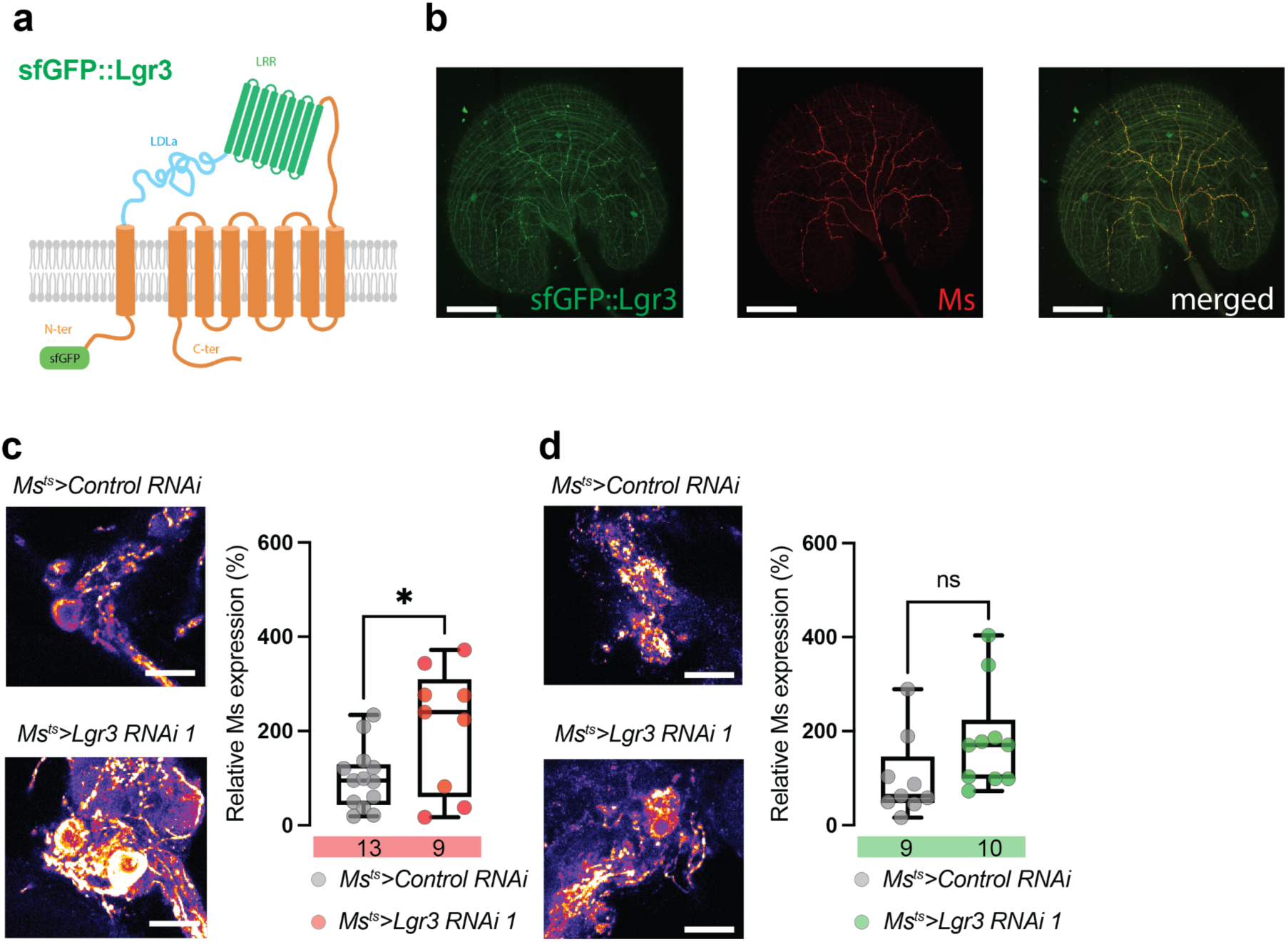
Visualization of Lgr3 protein and its role in regulating Ms peptide expression in the HCG. **(a)** Schematic of the sfGFP::Lgr3 protein construct, showing superfolder GFP (sfGFP) fused to the N-terminus of Lgr3. The Lgr3 receptor contains extracellular leucine-rich repeats (LRR), an LDLa domain, and seven transmembrane domains. **(b)** Confocal images of the crop terminals showing sfGFP::Lgr3 (green), Ms neurons (red), and merged panel. Lgr3 protein localises to Ms neuronal terminals in the crop, confirming expression at the protein level. Scale bars: 50 μm. **(c–e)** Confocal images for Ms peptide expression in the HCG of mated females or virgin females with or without *Lgr3* knockdown in Ms neurons. (c) *Ms^ts^>Control RNAi* vs. *Ms^ts^>Lgr3 RNAi 1* shows higher Ms expression upon *Lgr3* knockdown (*p < 0.05). (d) and **(e)** show no significant effect (ns) in virgin females. Scale bars: 20 μm.

**Extended Data Fig. 5.**
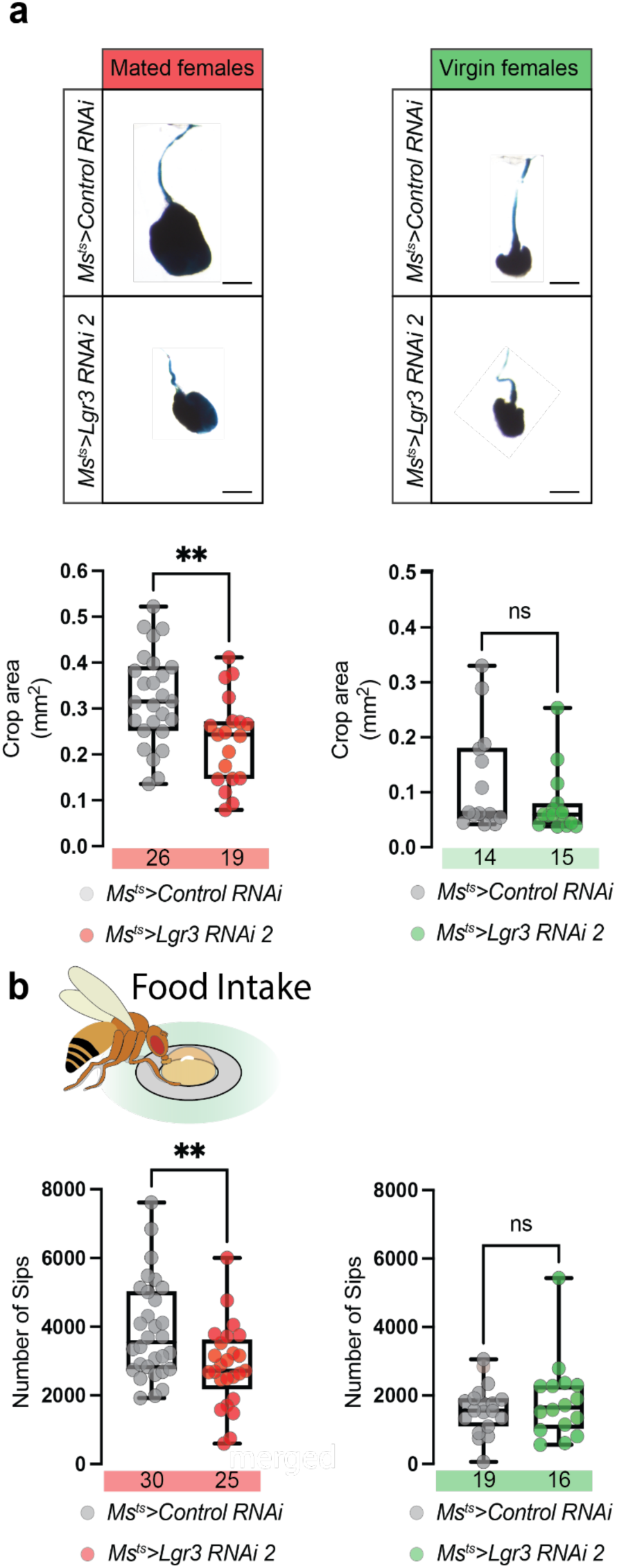
Independent *Lgr3* RNAi line confirms female-specific control of crop plasticity and food intake. **(a)** Representative images of dissected crops (top) and quantification of crop area (bottom) in mated females (red) and virgin females (green) expressing a second *Lgr3* RNAi line (*Lgr3 RNAi 2*) in Ms neurons. Knockdown of *Lgr3* significantly reduces crop size in mated females (**p < 0.01), but has no effect in virgin females. Scale bars: 200 μm. **(b)** Quantification of food intake using FlyPAD. *Lgr3* knockdown with Lgr3 RNAi 2 significantly reduces the number of sips in mated females (**p < 0.01), while virgin females show no significant difference.

**Extended Data Fig. 6.**
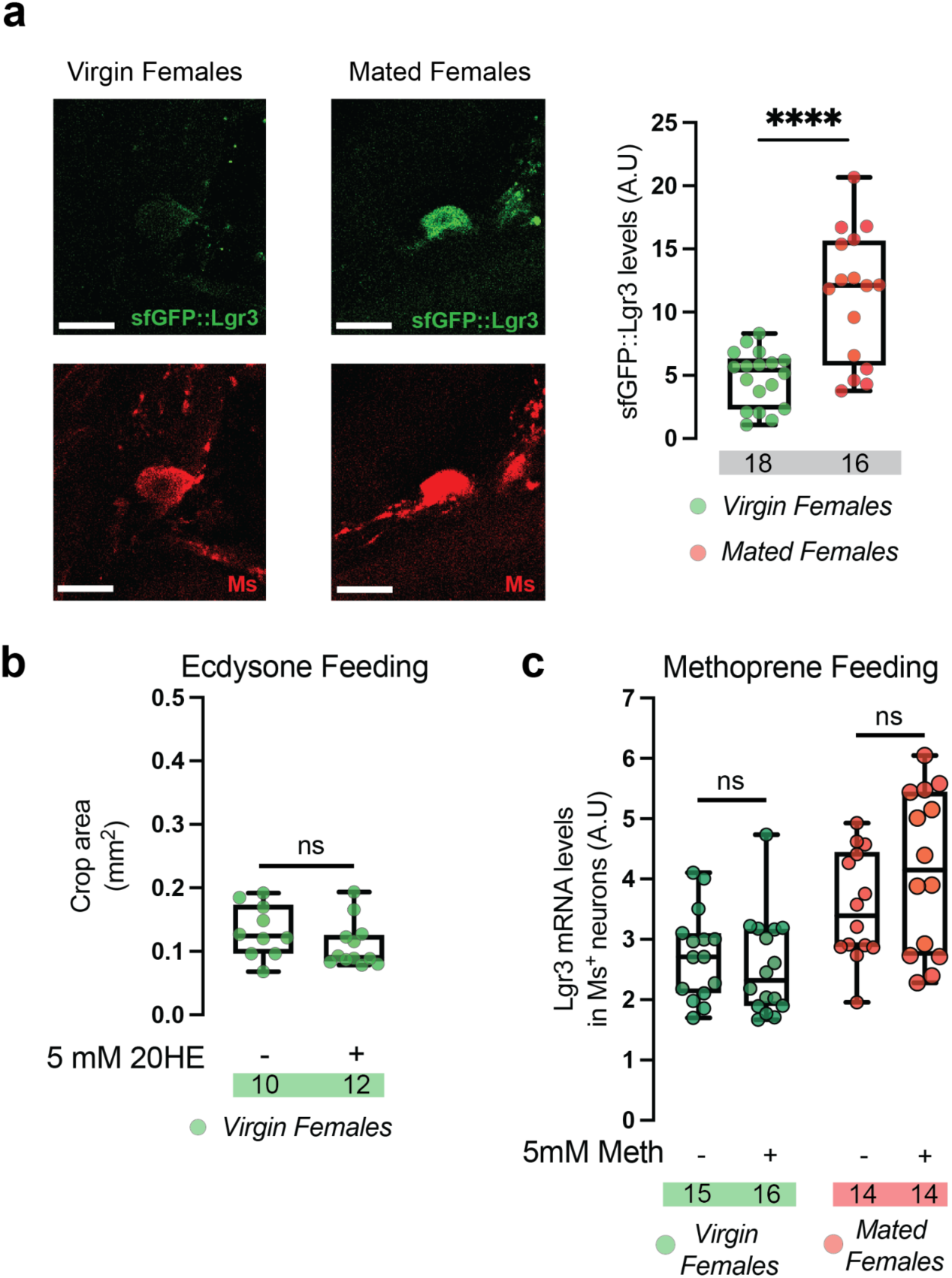
Lgr3 protein expression in the Ms neurons increases upon mating and is not induced by a juvenile hormone analogue. **(a)** Left: Confocal images of sfGFP::Lgr3 protein reporter (green) and Ms peptide (red) in virgin females and mated females. Lgr3 protein levels increase in mated females. Scale bars: 10 μm. Right: Quantification of sfGFP fluorescence intensity (A.U.) in Ms neurons of the HCG shows significantly higher Lgr3 protein levels in mated females compared to virgin females (****p < 0.0001). **(b)** Crop area quantification in virgin females fed with 5 mM 20-hydroxyecdysone (20HE). No significant effect is observed (ns), suggesting that ecdysone alone is not sufficient to induce gut remodelling. **(c)** Quantification of *Lgr3* mRNA levels by in situ hybridization in Ms neurons of virgin females, mated females, and virgin males after feeding with 5 mM methoprene (Meth), a juvenile hormone analogue. No significant changes in *Lgr3* expression are observed in any group (ns), indicating methoprene does not regulate *Lgr3* levels.

**Extended Data Fig. 7.**
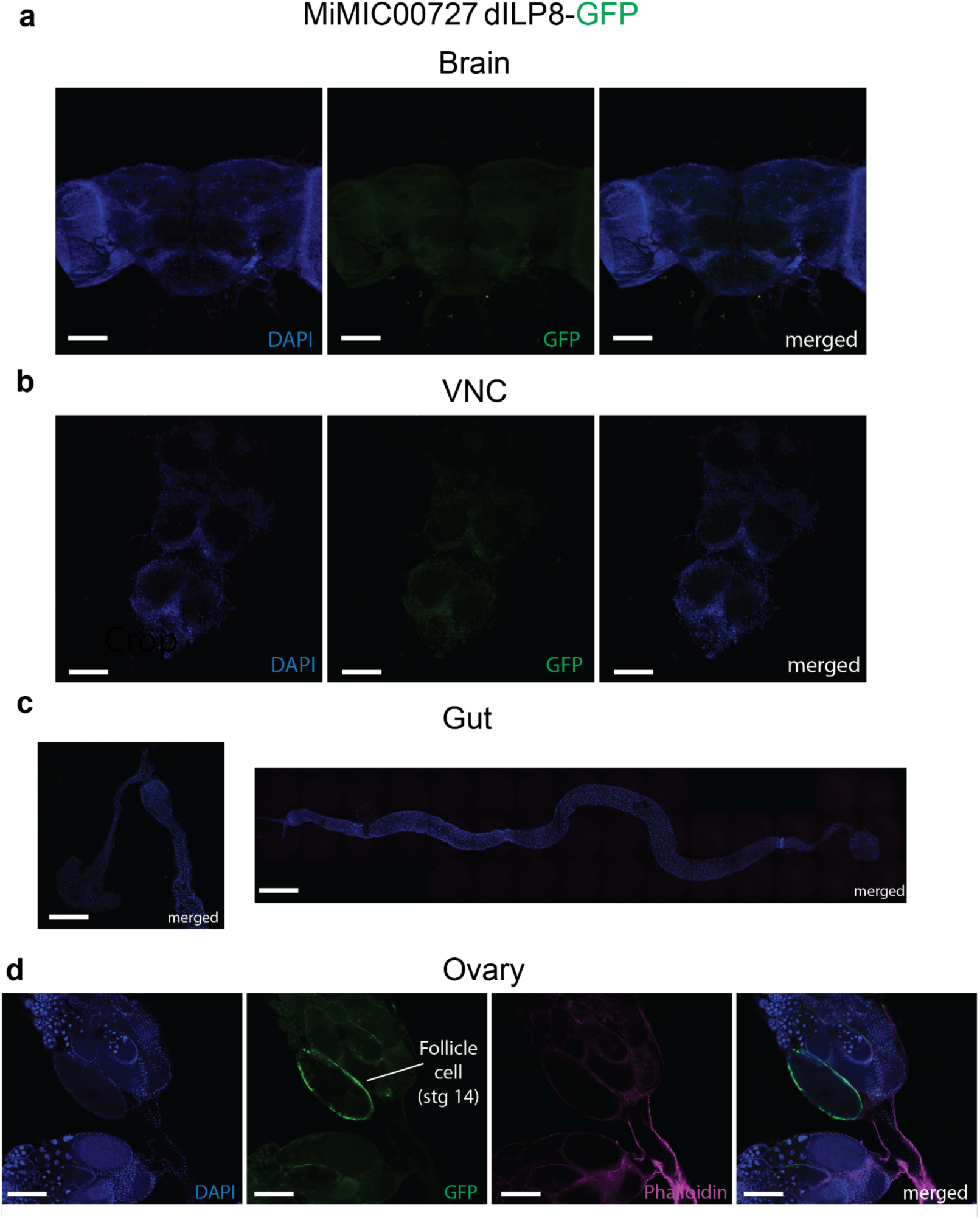
dILP8-GFP expression is restricted to late-stage follicle cells in the ovary. **(a–c)** Confocal images of adult female tissues from the *MiMIC00727 dILP8-GFP* reporter line, stained with DAPI (blue) and GFP (green). (a) Brain: No detectable GFP signal in the central brain region. (b) Ventral nerve cord (VNC): No detectable GFP expression. (c) Gut and crop: No GFP signal observed throughout the digestive tract. **(d)** Ovary: GFP signal is specifically detected in follicle cells surrounding stage 14 oocytes (green), confirming that *dILP8* is expressed in mature ovarian follicles. Phalloidin staining (magenta) outlines cell membranes. Scale bars: 50 μm (a–c), 20 μm (d).

**Extended Data Fig. 8.**
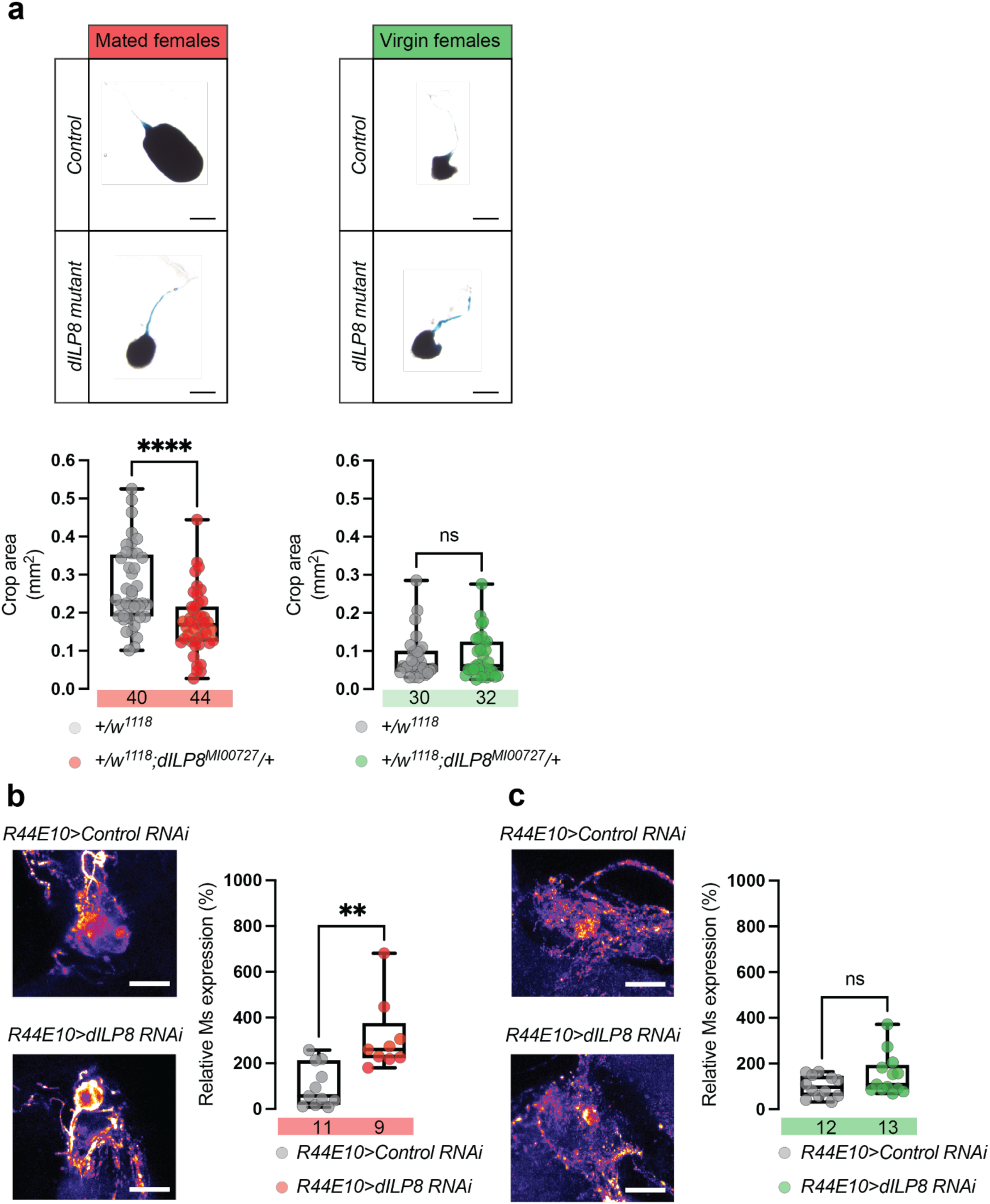
Loss of dILP8 impairs crop plasticity and food intake in a mating-dependent manner. **(a)** Representative images (top) and quantification (bottom) of crop size in control and *dILP8* mutant flies. In mated females (red), *dILP8* mutants (dilp8^MI00727^) show a significant reduction in crop area (****p < 0.0001), while no difference is seen in virgin females (green). Scale bars: 200 μm. **(b–c)** Confocal images of Ms peptide expression in the HCG. (b) *dILP8* RNAi driven by *R44E10-Gal4* significantly increases Ms peptide expression in mated females (**p < 0.001). (c) No effect is observed in virgin females (ns). Scale bars: 20 μm.

